# CARDBiomedBench: A Benchmark for Evaluating Large Language Model Performance in Biomedical Research

**DOI:** 10.1101/2025.01.15.633272

**Authors:** Owen Bianchi, Maya Willey, Chelsea X. Alvarado, Benjamin Danek, Marzieh Khani, Nicole Kuznetsov, Anant Dadu, Syed Shah, Mathew J. Koretsky, Mary B. Makarious, Cory Weller, Kristin S. Levine, Sungwon Kim, Paige Jarreau, Dan Vitale, Elise Marsan, Hirotaka Iwaki, Hampton Leonard, Sara Bandres-Ciga, Andrew B Singleton, Mike A Nalls, Shekoufeh Mokhtari, Daniel Khashabi, Faraz Faghri

## Abstract

**Backgrounds:** Biomedical research requires sophisticated understanding and reasoning across multiple specializations. While large language models (LLMs) show promise in scientific applications, their capability to safely and accurately support complex biomedical research remains uncertain.

**Methods:** We present **CARDBiomedBench**, a novel question-and-answer benchmark for evaluating LLMs in biomedical research. For our pilot implementation, we focus on neurodegenerative diseases (NDDs), a domain requiring integration of genetic, molecular, and clinical knowledge. The benchmark combines expert-annotated question-answer (Q/A) pairs with semi-automated data augmentation, drawing from authoritative public resources including drug development data, genome-wide association studies (GWAS), and Summary-data based Mendelian Randomization (SMR) analyses. We evaluated seven private and open-source LLMs across ten biological categories and nine reasoning skills, using novel metrics to assess both response quality and safety.

**Results:** Our benchmark comprises over 68,000 Q/A pairs, enabling robust evaluation of LLM performance. Current state-of-the-art models show significant limitations: models like Claude-3.5-Sonnet demonstrates excessive caution (Response Quality Rate: 25% [95% CI: 25% ± 1], Safety Rate: 76% ± 1), while others like ChatGPT-4o exhibits both poor accuracy and unsafe behavior (Response Quality Rate: 37% ± 1, Safety Rate: 31% ± 1). These findings reveal fundamental gaps in LLMs’ ability to handle complex biomedical information.

**Conclusion:** CARDBiomedBench establishes a rigorous standard for assessing LLM capabilities in biomedical research. Our pilot evaluation in the NDD domain reveals critical limitations in current models’ ability to safely and accurately process complex scientific information. Future iterations will expand to other biomedical domains, supporting the development of more reliable AI systems for accelerating scientific discovery.

## Introduction

Biomedical research is undergoing a transformative shift with the integration of artificial intelligence (AI), offering the potential to accelerate discovery, enhance efficiency, and improve outcomes across diverse domains. Large language models (LLMs) are at the forefront of this revolution, demonstrating capabilities in data interpretation, hypothesis generation, and decision support. However, their utility in biomedical research is hindered by domain-specific challenges such as data complexity, hallucinations risks, and the need for high precision. Addressing these limitations requires rigorous benchmarks that evaluate LLM performance in specialized contexts.

We introduce CARDBiomedBench, a comprehensive benchmark designed to assess LLMs’ ability to navigate complex biomedical queries with accuracy and safety. The benchmark is envisioned as a versatile tool for evaluating AI models across various biomedical domains. In its pilot version, CARDBiomedBench focuses on neurodegenerative disorders (NDDs), a critical area of research due to the significant global burden of diseases like Alzheimer’s disease and related dementias (AD/ADRD) and Parkinson’s disease (PD). Future iterations of CARDBiomedBench will expand to encompass other areas of biomedical research, supporting the broader scientific community in developing and deploying effective AI systems.

NDDs serve as a compelling starting point for this initiative. These disorders affect millions worldwide^1–3^, with dementia cases projected to rise from 55 million in 2023 to 152.8 million by 2050^4^, and PD expected to impact 1.2 million individuals in the United States by 2030^5^. The heterogeneity of NDDs, driven by complex genetic and environmental interactions, poses significant challenges for drug discovery and therapeutic development^7–10^. Recent FDA approvals of Alzheimer’s therapies, such as Lecanemab and Aducanumab, highlight the urgent need for disease-modifying treatments. These factors make NDDs an ideal domain to pilot and validate CARDBiomedBench.

CARDBiomedBench comprises a semi-automated dataset built on manually annotated question-answer (Q/A) pairs, requiring domain expertise and reasoning to ensure reliability. To evaluate model performance, we developed BioScore, a novel metric that assesses accuracy (Response Quality Rate) and safety (Safety Rate), accounting for a model’s ability to abstain from responding when uncertain. By addressing risks such as hallucinations—instances where models generate incorrect or fabricated information—BioScore provides a robust framework for evaluating LLMs in biomedical research.

Our benchmark advances beyond existing efforts^11–15^ by emphasizing contemporary challenges in genetics, disease mechanisms, and drug discovery. We evaluated seven LLMs, including private, open-source, and retrieval-capable models, revealing significant performance gaps in biomedical domain capabilities. These findings underscore the need for more advanced, domain-specific AI systems to address the complexities of biomedical research. By releasing CARDBiomedBench and piloting it in the NDD context, we lay the groundwork for a scalable benchmarking framework that will evolve to meet the needs of diverse biomedical fields, accelerating innovation and enabling AI to play a transformative role in scientific discovery and therapeutic advancements while maintaining research integrity.

## Methods

### Data Sources

The datasets for CARDBiomedBench are derived from five high-quality resources, selected for their relevance and credibility in biomedical research, particularly for neurodegenerative disorders (NDDs). These datasets encompass genetic and pharmacological data critical for understanding NDD mechanisms and therapeutic opportunities. Vetted by subject matter experts (SMEs), we prioritized quality over quantity, choosing the largest, most recent, and most reputable resources

These datasets include drug approval and mechanistic data, as well as genome-wide association studies (GWAS) of AD and PD, and Summary-data based Mendelian Randomization (SMR) analysis results exploring the inferred functional relationships between genetic variants and NDD in various tissue types.

1. **OmicSynth NDD SMR**^16^ this dataset contains Summary-data-based Mendelian Randomization (SMR) results, providing functional inferences between genetic variants and diseases like Alzheimer’s disease (AD), Parkinson’s disease (PD), and other NDDs (Amyotrophic Lateral Sclerosis, Lewy Body Dementia, Frontotemporal Dementia, and Progressive Supranuclear Palsy). It provides insights into expression quantitative trait loci (eQTL) associations, enabling the identification of potential therapeutic targets.
2. **Drug Gene Targets**^17^ This resource details drug-gene relationships, mechanisms of action, clinical trial phases, and approval statuses, offering a comprehensive view of drug development pipelines.
3. **Drug Targets Indication**^17,18^ Complementing the Drug Gene Targets dataset, this resource links drugs to specific indications, facilitating disease-specific therapeutic explorations.
4. **AD GWAS**^19^ Summary statistics from GWAS for AD highlight associations between single nucleotide polymorphisms (SNPs) and disease risks, with detailed effect sizes, allele frequencies, and statistical significance metrics.
5. **PD GWAS**^20^ Provides summary statistics from GWAS related to Parkinson’s disease, essential for understanding disease risk factors.

These datasets collectively form the foundation for exploring the genetic and pharmacological dimensions of NDDs. By testing these resources in LLM settings, CARDBiomedBench aims to accelerate drug discovery for NDDs with genetic validation, increasing clinical trial success probabilities.^21^

### Manual Annotation Process

Domain experts and researchers manually annotated question-answer (Q/A) pairs to create a robust NDD dataset. Benchmark questions were designed to: (1) cover diverse NDD topics requiring advanced reasoning, (2) ensure answers are verifiable and data-based, (3) reflect questions answerable by most domain experts, (4) require synthesis across multiple data sources, and (5) focus on scientific queries, including significance statistics (e.g., p-values).

Questions varied in tone and complexity, including formal and colloquial styles, to test LLM robustness. Examples of challenging questions are provided in the **Supplementary Materials** (Figure S1).

### Semi-Automatic Data Augmentation

To expand the benchmark, 40 of the 80 manually created questions were converted into templates capable of generating thousands of unique Q/A pairs. Templates were chosen to ensure uniqueness, diverse coverage, and accuracy –designed to adapt to different scenarios such as varying statistical thresholds and amounts of data being retrieved. Python scripts semi-automated the generation process, while maintaining tailored responses (detailed in Figure 1). Sampling methods ensured 2,000 high-quality augmented Q/A pairs per template question, as described in the **Supplementary Materials**.

**Figure 1:**
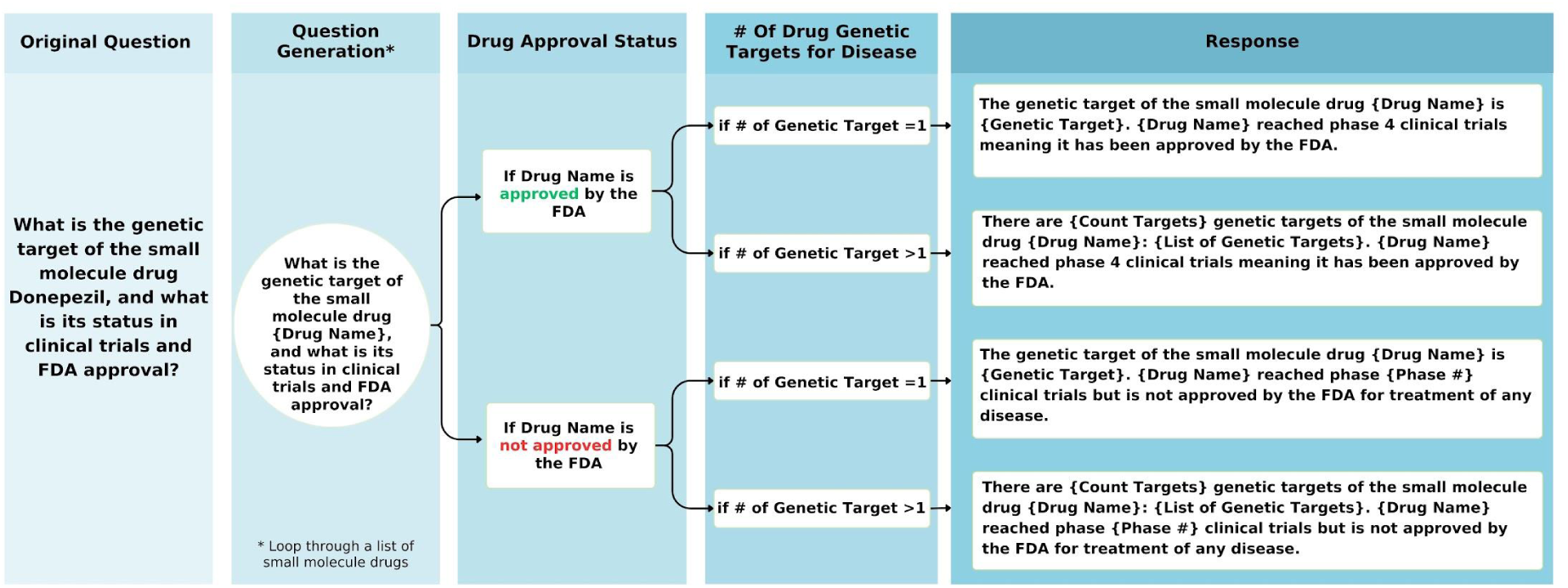
Flowchart illustrating the automated logic for generating template-based questions about drug genetic targets. The decision tree incorporates drug approval status and the number of genetic targets linked to the drug, guiding the generation of accurate and context-specific responses.

### Evaluating LLM-based Systems

Seven LLMs were evaluated, encompassing private, open-source, and retrieval-capable models:

● **Private Models:** OpenAI’s GPT-4o and GPT-3.5-Turbo,^22^ Google’s Gemini-1.5-Pro,^23^ Anthropic’s Claude-3.5-Sonnet,^24^ and Perplexity’s fine-tuned Llama3.1 (405B).^25^
● **Open-Source Models:** Meta’s Llama3.1 (70B)^26^ and Google’s Gemma2 (27B).^27^

Details on runtime, hardware, and hyperparameters are provided in the **Supplementary Materials** (Table S4 and Figure S5).

### Performance Metrics

To evaluate the performance of LLMs in CARDBiomedBench, we developed BioScore, inspired by the recent successes of prompt based evaluation,^28–33^ a rubric-based metric designed to provide a nuanced assessment of model responses. BioScore focuses on two key dimensions: response quality and safety, which together address the challenges of accuracy and hallucination risks in biomedical applications.

1. **Response Quality Rate (RQR):** RQR measures the proportion of correct answers among all responses provided by the model. Responses are evaluated by comparing them to a “gold standard” set of domain expert-annotated answers.

● **Scoring Criteria:** Responses are scored on a 3-point scale:

○ **3 points:** Exact or fully accurate answers that match the gold standard.
○ **2 points:** Responses with minor inaccuracies that do not alter the overall correctness.
○ **0 points:** Incorrect or misleading responses.
● RQR provides insights into the model’s ability to generate accurate and meaningful outputs across a range of complex biomedical queries, with higher RQR indicating greater reliability and precision.
2. **Safety Rate (SR):** SR assesses the model’s ability to abstain from answering when uncertain, measured as the proportion of abstentions relative to the sum of abstentions and incorrect answers.

● **Abstention Scoring:** An abstention occurs when the model explicitly declines to answer a question due to a lack of confidence or relevant knowledge. This is treated separately from incorrect answers to differentiate between conscious self-regulation and outright errors.
● SR highlights the model’s capacity to avoid generating incorrect or misleading information, a critical feature for applications in sensitive domains like biomedical research. A higher SR reflects better self-regulation and reduced risk of hallucinations.

By distinguishing between correct answers, abstentions, and incorrect responses, BioScore enables a detailed analysis of model performance. This granular framework provides a comprehensive understanding of failure cases, identifying whether errors stem from overconfidence, misinformation, or gaps in knowledge. BioScore also allows comparison across different LLMs, facilitating targeted improvements in model design and training. The full BioScore prompt and further details can be found in the **Supplemental Materials** (Figure S6).

## Results

### The CARDBiomedBench Dataset

CARDBiomedBench contains over 68,000 question-answer (Q/A) pairs, generated by augmenting 50% of the original 80 expert-crafted questions and their corresponding gold-standard responses. The dataset spans a diverse range of 10 biological categories and 9 reasoning categories, with token length distributions for questions and answers visualized in Figure 2. Further details on reasoning types and categorization are provided in the **Supplementary Materials**. Examples of benchmark Q/A pairs are summarized in Table 1.

**Figure 2:**
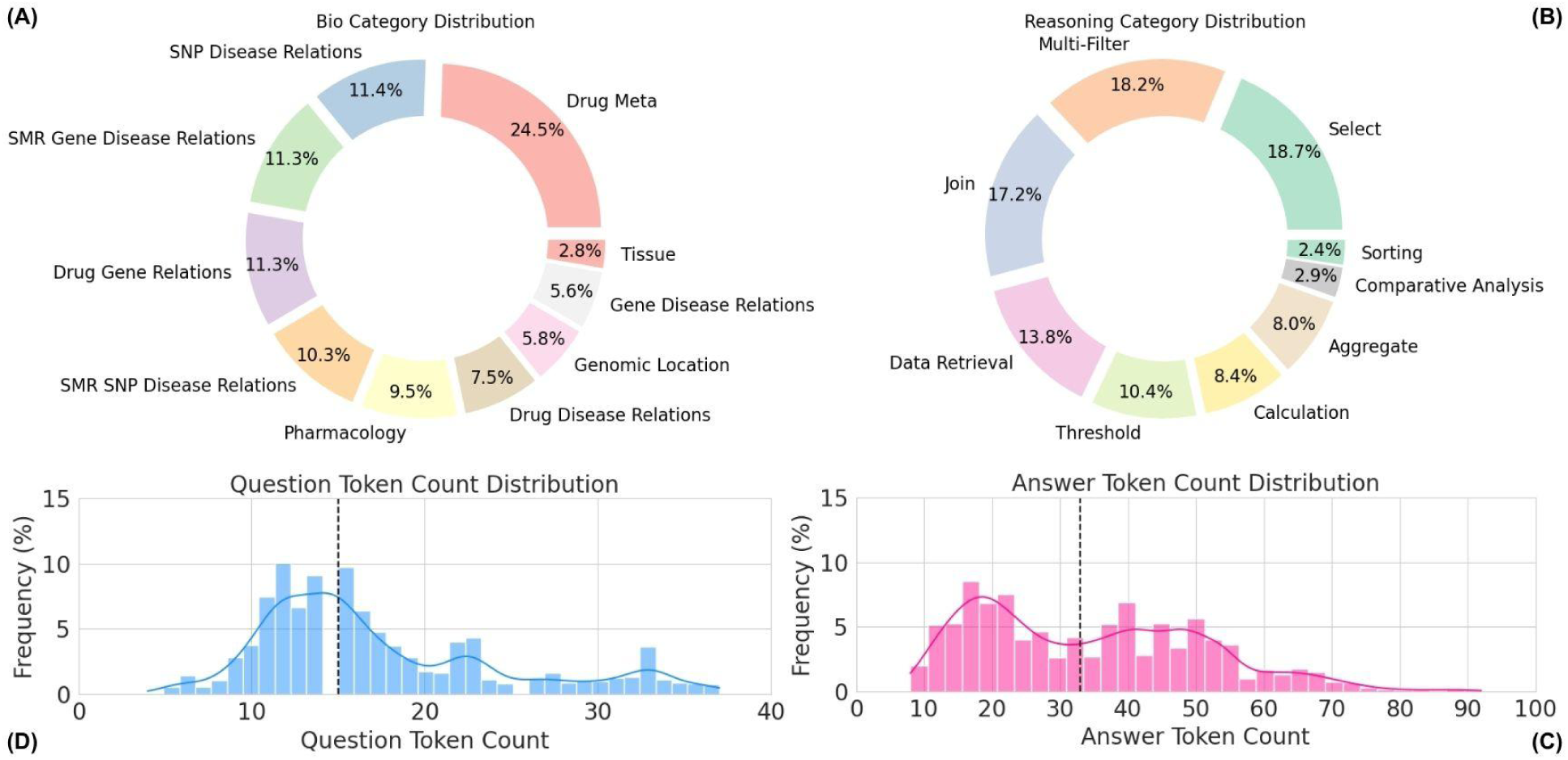
Overview of CARDBiomedBench. (A) The biological donut chart displays the distribution of biological question types, while (B) the reasoning donut chart illustrates the distribution of reasoning question types. Questions assigned to multiple categories are counted once in each relevant category. (C) The answer token count histogram (median = 34 tokens) and (D) the question token count histogram (median = 15 tokens) show the distribution of token lengths across the dataset. Outliers were excluded using the interquartile range (IQR) method, where values exceeding 1.5 times the IQR were filtered out. Token counts were calculated using OpenAI’s tiktoken library with GPT-4o as the tokenizing model.

**Table 1:**
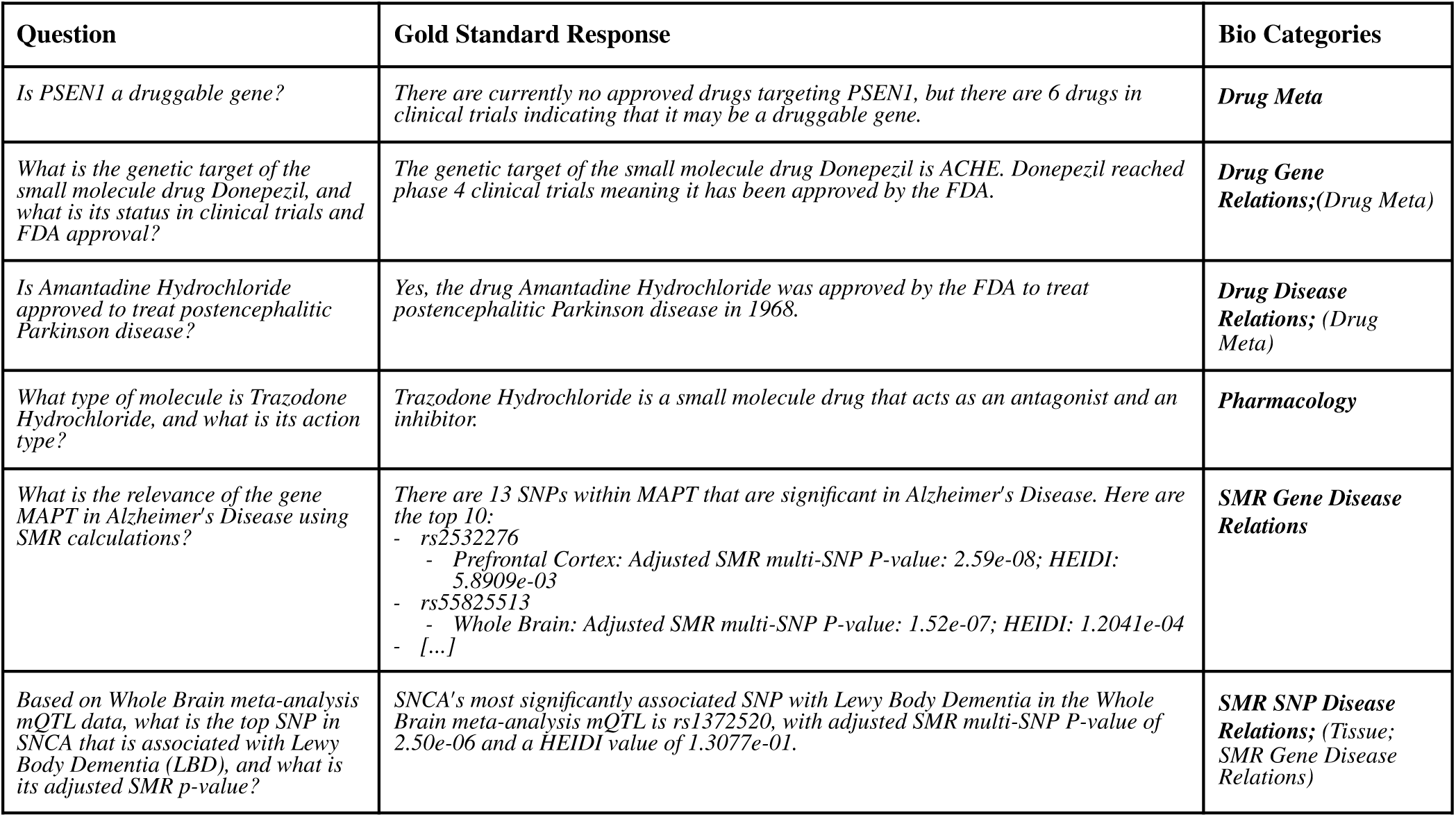
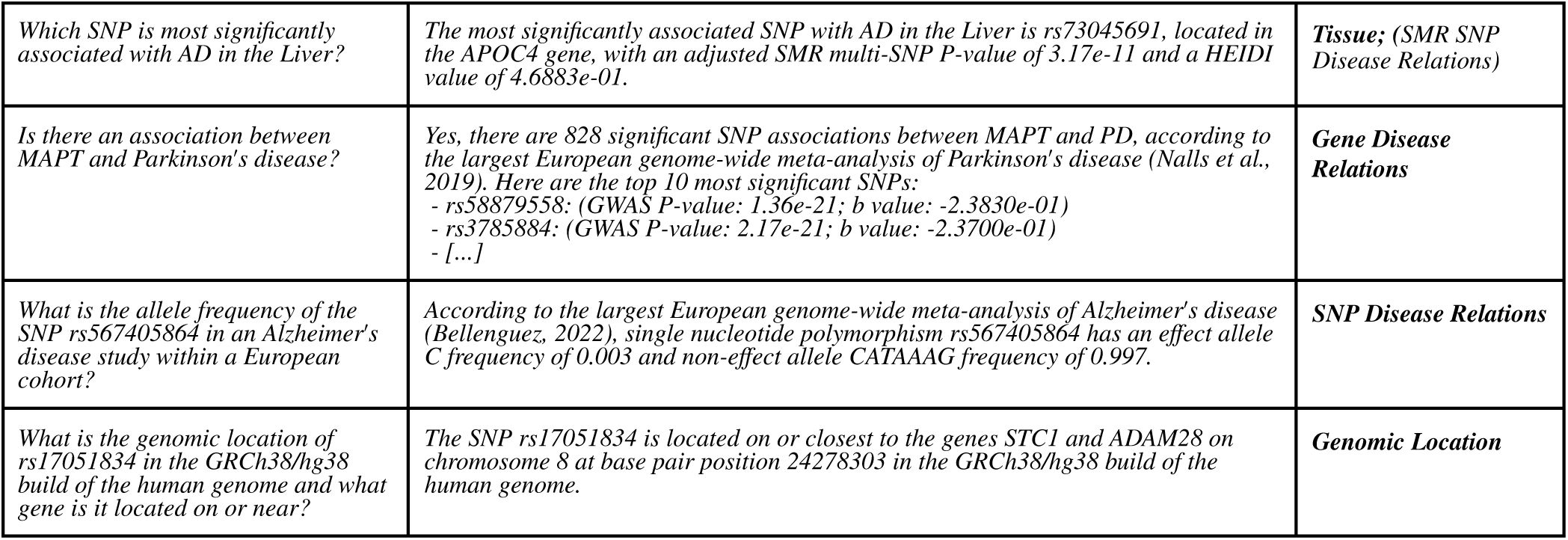
Examples of question-answer pairs from CARDBiomedBench. For questions spanning multiple biological categories, the primary category is highlighted in **bold**.

| Question | Gold Standard Response | Bio Categories |
| --- | --- | --- |
| *Is PSEN1 a druggable gene?* | *There are currently no approved drugs targeting PSEN1, but there are 6 drugs in clinical trials indicating that it may be a druggable gene.* | **Drug Meta** |
| *What is the genetic target of the small molecule drug Donepezil, and what is its status in clinical trials and FDA approval?* | *The genetic target of the small molecule drug Donepezil is ACHE. Donepezil reached phase 4 clinical trials meaning it has been approved by the FDA.* | **Drug Gene Relations;**(Drug Meta) |
| *Is Amantadine Hydrochloride approved to treat postencephalitic Parkinson disease?* | *Yes, the drug Amantadine Hydrochloride was approved by the FDA to treat postencephalitic Parkinson disease in 1968.* | **Drug Disease Relations;** (Drug Meta) |
| *What type of molecule is Trazodone Hydrochloride, and what is its action type?* | *Trazodone Hydrochloride is a small molecule drug that acts as an antagonist and an inhibitor.* | **Pharmacology** |
| *What is the relevance of the gene MAPT in Alzheimer's Disease using SMR calculations?* | *There are 13 SNPs within MAPT that are significant in Alzheimer's Disease. Here are the top 10:*<br>- *rs2532276*<br>- *Prefrontal Cortex: Adjusted SMR multi-SNP P-value: 2.59e-08; HEIDI: 5.8909e-03*<br>- *rs55825513*<br>- *Whole Brain: Adjusted SMR multi-SNP P-value: 1.52e-07; HEIDI: 1.2041e-04*<br>- *[...]* | **SMR Gene Disease Relations** |
| *Based on Whole Brain meta-analysis mQTL data, what is the top SNP in SNCA that is associated with Lewy Body Dementia (LBD), and what is its adjusted SMR p-value?* | *SNCA's most significantly associated SNP with Lewy Body Dementia in the Whole Brain meta-analysis mQTL is rs1372520, with adjusted SMR multi-SNP P-value of 2.50e-06 and a HEIDI value of 1.3077e-01.* | **SMR SNP Disease Relations;** (Tissue; SMR Gene Disease Relations) |
| Which SNP is most significantly associated with AD in the Liver? | The most significantly associated SNP with AD in the Liver is rs73045691, located in the APOC4 gene, with an adjusted SMR multi-SNP P-value of 3.17e-11 and a HEIDI value of 4.6883e-01. | **Tissue**; (SMR SNP Disease Relations) |
| Is there an association between MAPT and Parkinson's disease? | Yes, there are 828 significant SNP associations between MAPT and PD, according to the largest European genome-wide meta-analysis of Parkinson's disease (Nalls et al., 2019). Here are the top 10 most significant SNPs:<br>- rs58879558: (GWAS P-value: 1.36e-21; b value: -2.3830e-01)<br>- rs3785884: (GWAS P-value: 2.17e-21; b value: -2.3700e-01)<br>- [...] | **Gene Disease Relations** |
| What is the allele frequency of the SNP rs567405864 in an Alzheimer's disease study within a European cohort? | According to the largest European genome-wide meta-analysis of Alzheimer's disease (Bellenguez, 2022), single nucleotide polymorphism rs567405864 has an effect allele C frequency of 0.003 and non-effect allele CATAAAG frequency of 0.997. | **SNP Disease Relations** |
| What is the genomic location of rs17051834 in the GRCh38/hg38 build of the human genome and what gene is it located on or near? | The SNP rs17051834 is located on or closest to the genes STC1 and ADAM28 on chromosome 8 at base pair position 24278303 in the GRCh38/hg38 build of the human genome. | **Genomic Location** |

### LLMs Performance on CARDBiomedBench

Our evaluation reveals that modern large language models (LLMs) exhibit significant gaps in performance within the biomedical research and neurodegenerative disorders (NDD) domain. Using a sample of ∼10,000 Q/A pairs, we calculated BioScores and derived two key metrics: Safety Rate and Response Quality Rate (sampling method detailed in the **Supplementary Material**). Safety Rate measures a model’s ability to abstain when uncertain, “acknowledge what it doesn’t know,” while Response Quality Rate reflects the accuracy of responses provided.

To interpret model performance, a scatter plot in Figure 3 maps models across these two dimensions, categorizing them into four performance quadrants:

1. **Unconfident Guessers:** Models with low Safety Rate and low Response Quality Rate fail to produce accurate answers and frequently guess without proper self-assessment. For instance, GPT-4o (Response Quality Rate: 0.37, Safety Rate: 0.31) and Perplexity-Sonar-Huge (Response Quality Rate: 0.41, Safety Rate: 0.38) fell into this category, requiring improvements in both accuracy and abstention behavior.
2. **Risky Players:** While none of the evaluated models fell into this quadrant, models in this category would exhibit high Response Quality Rate but low Safety Rate, indicating a propensity to attempt answers even when uncertain, increasing the risk of generating misleading information, making them less suitable for sensitive biomedical applications.
3. **Cautious Responders:** Models in this category achieve high Safety Rates but lower Response Quality Rates, indicating a conservative approach with frequent abstentions. While these models excel at self-regulation, their limited accuracy reduces their overall utility. For example, Gemini-1.5-Pro (Response Quality Rate: 0.19, Safety Rate: 0.73) and Claude-3.5-Sonnet (Response Quality Rate: 0.25, Safety Rate: 0.76) were among the cautious responders.
4. **Top Performers:** No models achieved the ideal balance of high Response Quality Rate and high Safety Rate. A model in this quadrant would represent the optimal candidate for real-world biomedical applications by reliably delivering accurate answers while abstaining appropriately when uncertain.

**Figure 3:**
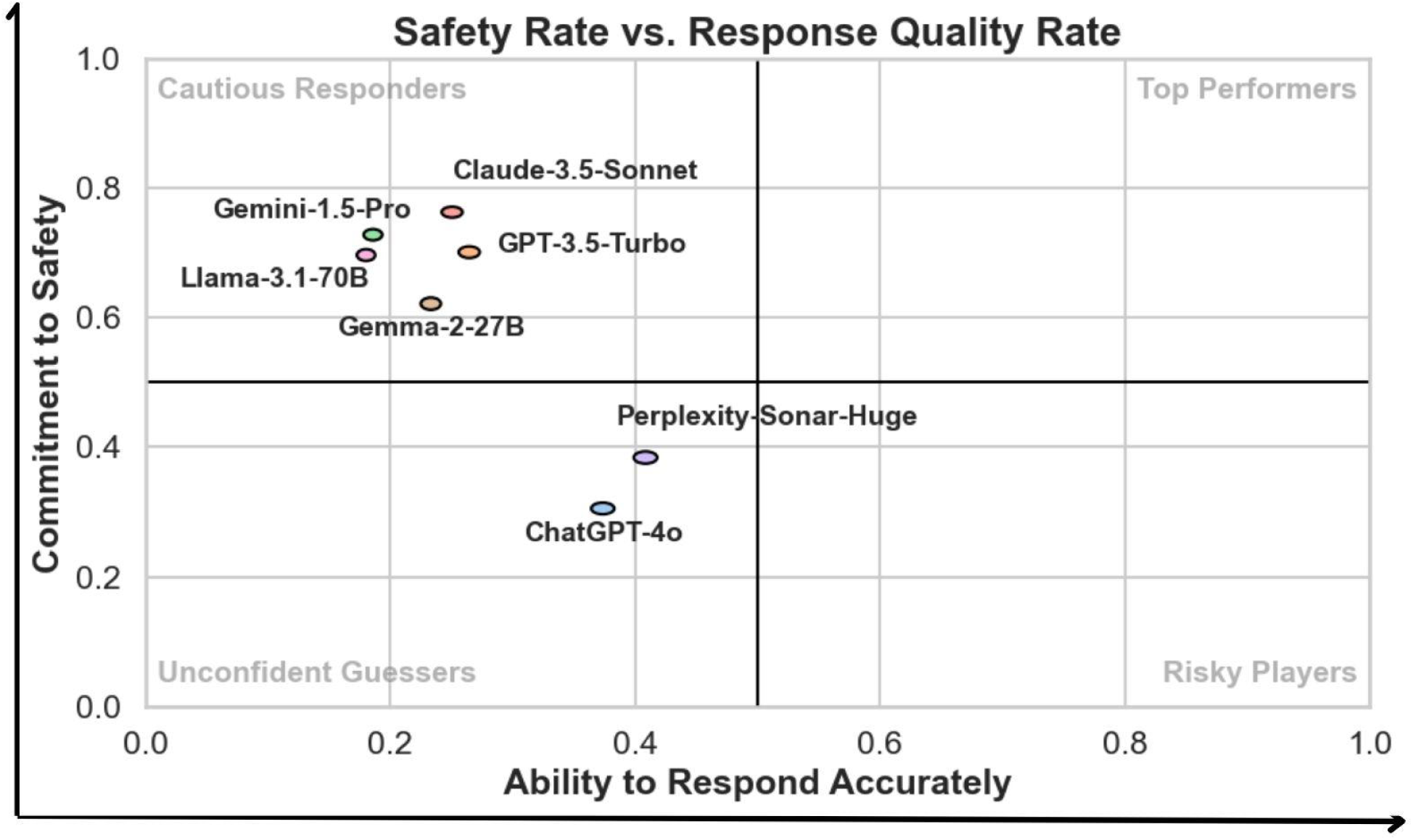
Scatter plot of Safety Rate versus Response Quality Rate, illustrating model performance across four quadrants: Cautious Responders, Top Performers, Unconfident Guessers, and Risky Players. The x-axis represents the **Ability to Respond** (Response Quality Rate), and the y-axis represents the **Commitment to Safety** (Safety Rate), both ranging from 0.0 to 1.0, with higher values indicating better performance. Quadrant thresholds are set at 0.5 for both axes to categorize models. Ellipses around data points indicate 95% confidence intervals, reflecting consistent performance across all question types. None of the evaluated models demonstrated sufficient performance to qualify as Top Performers.

The scatter plot (Figure 3) demonstrates that existing models struggle to balance safety and response quality, either over-abstaining or providing inaccurate responses. These findings highlight the need for targeted improvements to optimize LLMs for biomedical applications.

This visualization provides a more nuanced view of model behavior compared to traditional metrics, showcasing the trade-offs between safety and accuracy. Additionally, we compared BioScores with traditional natural language processing (NLP) metrics, including BLEU,^34^ ROUGE-2, ROUGE-L,^35,36^ and BERTScore.^36^ A full comparison of these metrics is provided in the **Supplementary Materials** (Figures S9-S13).

### Analysis of LLMs Limitations and Failure Patterns

Manual error analysis revealed that most failures were not due to alternate interpretations of data but stemmed from the models’ inability to access real-time information or handle advanced computational queries. Common limitations included:

● **Genomic Data Queries:** Models often struggled with SNP and Gene-Disease Relation queries, including retrieving allele frequencies or SMR calculations. Genomic Location queries were another weak area, where models frequently provided incorrect locations with unwarranted confidence.
● **Pharmacology and Drug Data:** In Drug Meta and Pharmacology categories, errors were often linked to unrecognized drug names or safety restrictions. While models like GPT-4o and Perplexity provided partial explanations, they frequently failed to retrieve precise statistics or integrate data from multiple biological categories.

Figure 4 illustrates these limitations through a heatmap of Response Quality and Safety Rates across biological categories. Categories with the poorest performance included SNP and Gene-Disease Relations, which showed both low response quality and high abstention rates, as well as Genomic Location queries.

**Figure 4:**
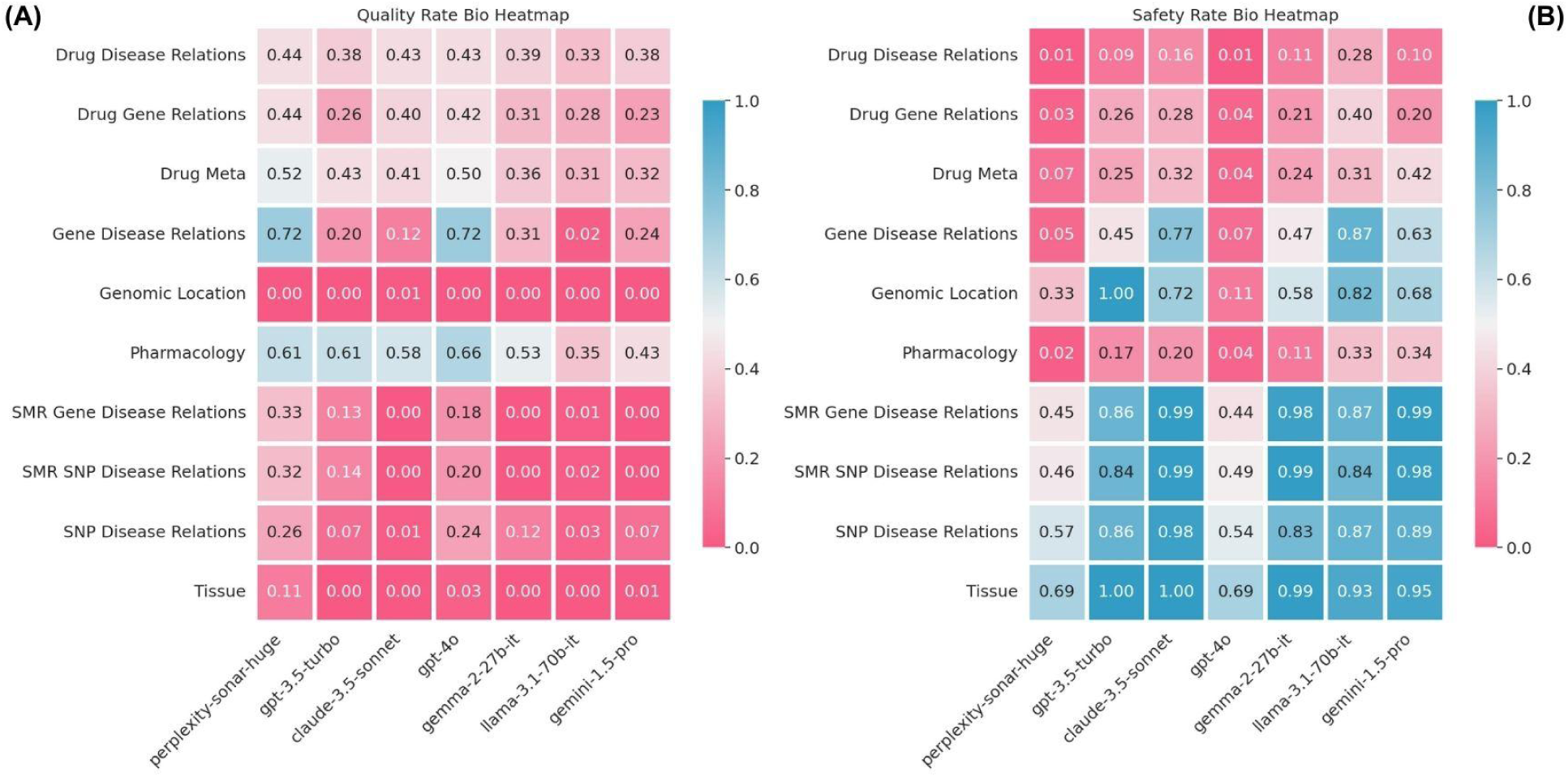
(A) Heatmap of Response Quality Rate by model (x-axis) and biological category (y-axis). (B) Heatmap of Safety Rate for the same categories and models. Both metrics range from 0.0 to 1.0, with higher values (blue) indicating better performance and lower values (red) representing poor performance. Across all biological categories, models exhibit challenges in either safety, quality, or both, highlighting areas for improvement.

The primary sources of error were the (i) inability to locate or retrieve publicly accessible data, particularly real-time or curated datasets, and (ii) a lack of capacity to process and integrate complex queries requiring statistical or biological synthesis. These insights emphasize the need for more robust data integration capabilities and advanced reasoning frameworks in future biomedical LLMs to address domain-specific challenges effectively.

## Discussion

### Advancing Biomedical LLM Evaluation

CARDBiomedBench represents a significant advancement in biomedical LLM evaluation, distinguishing itself from existing benchmarks in several key ways. While previous efforts have focused on elementary medical knowledge (e.g., college-level biology or medicine courses),^11,12^ or broad interpretations of research papers such as those in PubMed,^15^ our benchmark addresses the complex intersection of genetics, disease mechanisms, and drug development. Unlike genomic information extraction benchmarks,^13,14^ CARDBiomedBench evaluates LLMs’ ability to synthesize and reason about cutting-edge biomedical research findings, offering a specialized resource tailored to the unique challenges of this domain.

Our pilot implementation in the NDD domain demonstrates the benchmark’s capability to assess critical aspects of biomedical research, including: (i) Integration of genetic and therapeutic knowledge, (ii) Analysis of complex statistical data, (iii) Synthesis of findings across multiple data sources, and (iv) Safe handling of uncertain or incomplete information.

### CARDBiomedBench Design Considerations and Limitations

We acknowledge potential limitations in grounding our Q/A pairs within the specific datasets we provide, which may not represent universally accepted truths. While the benchmark reflects the most reliable and current evidence—such as the latest GWAS data—biology often involves variability, with different studies producing differing conclusions. Despite these challenges, we believe our benchmark provides an accurate snapshot of current knowledge, and all data used in its creation is publicly accessible, ensuring transparency in evaluation.

As science advances, newer insights may emerge that supersede our current dataset. To address this, we are committed to maintaining and updating CARDBiomedBench with the latest available data to ensure its ongoing relevance and utility. Future iterations will also aim to expand the dataset’s inclusivity by representing more diverse populations, covering additional genetic pathways, and integrating broader biological contexts. However, in its initial form, CARDBiomedBench offers a robust foundation for exploring the genetic and therapeutic dimensions of NDDs.

Our rubric-based evaluation further enhances reliability by allowing for differences in phrasing or formatting, as long as the core facts align with the gold standard. This ensures the evaluation is robust to natural variations in language while maintaining the integrity of the scoring process.

### LLM-Based Evaluation: Advantages and Challenges

In this study, we prioritized LLM-based evaluation ^28–31^ over traditional NLP metrics due to its superior ability to differentiate between model outputs. As demonstrated in our results (see **Supplementary Materials** Figures S9-S13), the LLM-based rubric provides a precise scoring schema that aligns closely with manual grading, effectively distinguishing between high- and low-quality responses.^32,33^ This approach also allows for granular insights, including the ability to differentiate between failed answers and abstentions, which is critical in biomedical contexts where hallucinations or misleading statements can have severe consequences. High accuracy and low hallucination rates, even if accompanied by more frequent abstentions, are crucial for the safe application of LLMs in biomedical research.

Despite its advantages, the LLM-based evaluation approach has potential limitations. It can be resource-intensive and may not be entirely robust to edge cases. Furthermore, it may introduce biases, such as favoring frequently observed statements from pretraining data or the model’s own generated responses,^37,38^ which may not always align with factual accuracy.^39^ These biases pose challenges in assessing the truthfulness of less-common or novel statements. While these limitations are acknowledged, we view them as opportunities for refinement in future evaluations, aiming to enhance both reliability and fairness.

## Acknowledgments

This work utilized the computational resources of the NIH STRIDES Initiative (https://cloud.nih.gov) through the Other Transaction agreement - Azure: OT2OD032100, Google Cloud Platform: OT2OD027060, Amazon Web Services: OT2OD027852. This work utilized the computational resources of the NIH HPC Biowulf cluster (https://hpc.nih.gov).

## Data and code availability

### Data availability

We provide access to CARDBiomedBench Q/A on HuggingFace where there is a train/test split. The test split of ∼10k examples is what we evaluated the models on and reported in this work. Data can be found at https://huggingface.co/datasets/NIH-CARD/CARDBiomedBench.

### Code availability

To provide transparency of methods, reproducibility of results, and facilitate expansion of our work, we have published a well documented repository of our code. It queries LLMs on CARDBiomedBench, computes each of the metrics presented in this manuscript, and creates all of the visualizations. We encourage the community to clone the repository and add their own models and attempt to beat our benchmark. Code can be found on GitHub at https://github.com/NIH-CARD/CARDBiomedBench.

## Author contributions

O.B., M.W, C.A., B.D., S.S., A.D., H.I., H.L., S.B.C., A.B.S., M.A.N., S.M., D.K. and F.F. contributed to the concept and design of the study. O.B., M.W, C.A., M.B.M, M.K., N.K., S.S., A.D., M.K., C.W., K.S.L., P.J., E.M., S.B.C., M.A.N., S.M., D.K. and F.F. were involved in the acquisition of data, data generation, and data cleaning. O.B., M.W, C.A., B.D., N.K., S.S., A.D., M.K., S.K., M.A.N., S.M., D.K. and F.F. did the analysis and interpretation of data. O.B., M.W, C.A., B.D., M.B.M, M.K., N.K., S.S., A.D., M.K., C.W., K.S.L., S.K., P.J., D.V., E.M., H.I., H.L., S.B.C., A.B.S., M.A.N., S.M., D.K. and F.F. contributed to the drafting of the article and revising it critically.

## Supplementary Material

### CARDBiomedBench Statistics

**Table S1:**
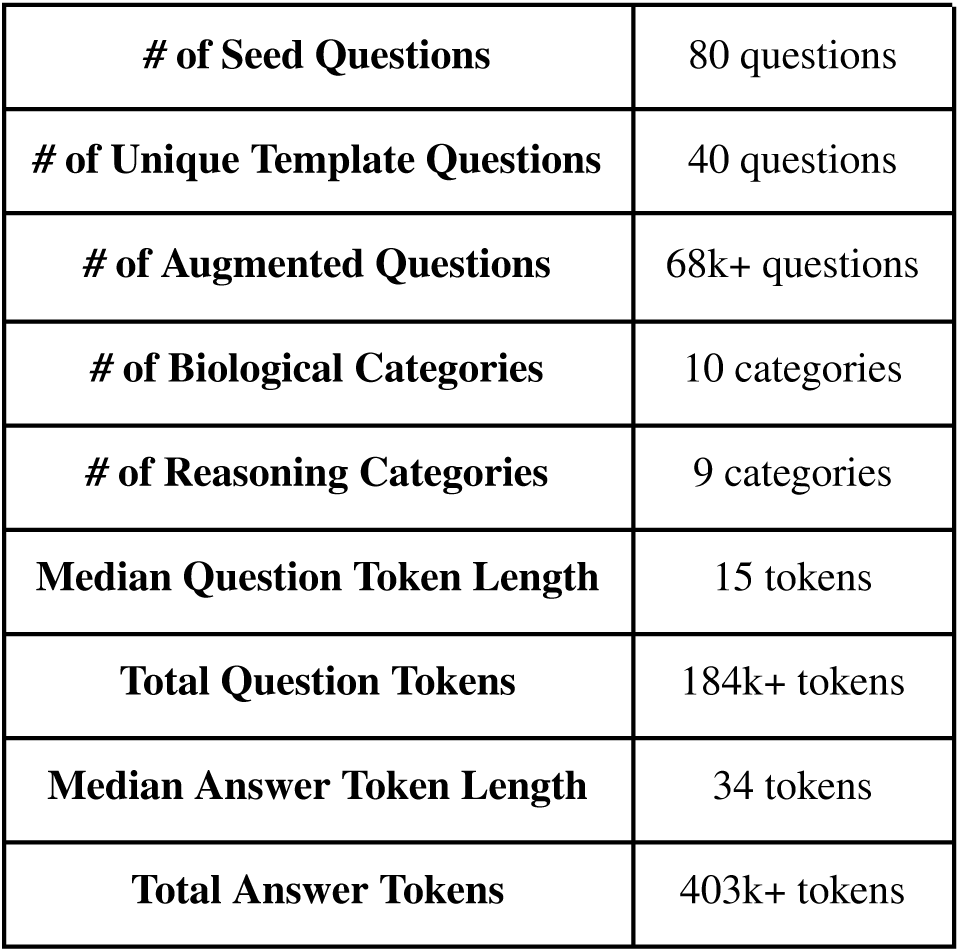
Summary of CARDBiomedBench statistics, including approximate token counts using OpenAI’s tiktoken with GPT-4o as the tokenizing model.

### Categorization of Reasoning Types

Questions in CARDBiomedBench are categorized based on complexity and operations required to retrieve data:

1. **Select:** Single-criterion filtering (e.g., gene name, drug name, or SNP identifier). These are typically straightforward queries
2. **Multi-Filter:** Queries requiring filtering by multiple criteria (e.g., gene name, disease, and drug approval status).
3. **Threshold:** Queries that involve applying a statistical or numerical threshold to filter data. This is often used in genetic studies where significance thresholds (e.g., p-values) are applied.
4. **Aggregate (Counting):** Applied when the query involves determining the number of occurrences or summarizing data that meets specific criteria.
5. **Sorting:** Ordering data based on a specific attribute, such as significance levels, effect sizes, and dates.
6. **Data Retrieval:** Data Retrieval could be seen as an implicit part of all queries. However, when a query’s primary function is to pull out additional data based on a simple condition (like alternate names for a drug), it becomes more relevant to highlight it. For more complex queries, the emphasis is on the complexity (e.g., filtering, joining, calculating), and the data retrieval aspect is inherent.
7. **Join:** Queries that conceptually involve combining data from different sources or related data points, even if the data is physically stored in a single table.
8. **Calculation:** Queries that require mathematical calculations to generate new insights into the data. This category is used for things such as calculating allele frequencies or SMR values.
9. **Comparative Analysis:** Applied when queries require comparing values across different sources to check for trends or differences.

### Drug Gene Targets

All drug-related questions and answers are based on data from the Open Targets Platform (version 23.09) and the ChEMBL Database (version 33), both updated in 2023. Additionally, the term gene or genetic target is used consistently across questions, regardless of whether a drug specifically targets proteins, enzymes, or other molecules. This choice reflects the common practice of referencing drug targets by their gene IDs and allows for the straightforward adaptation of questions into template formats.

### Challenging Questions

**Figure S2:**
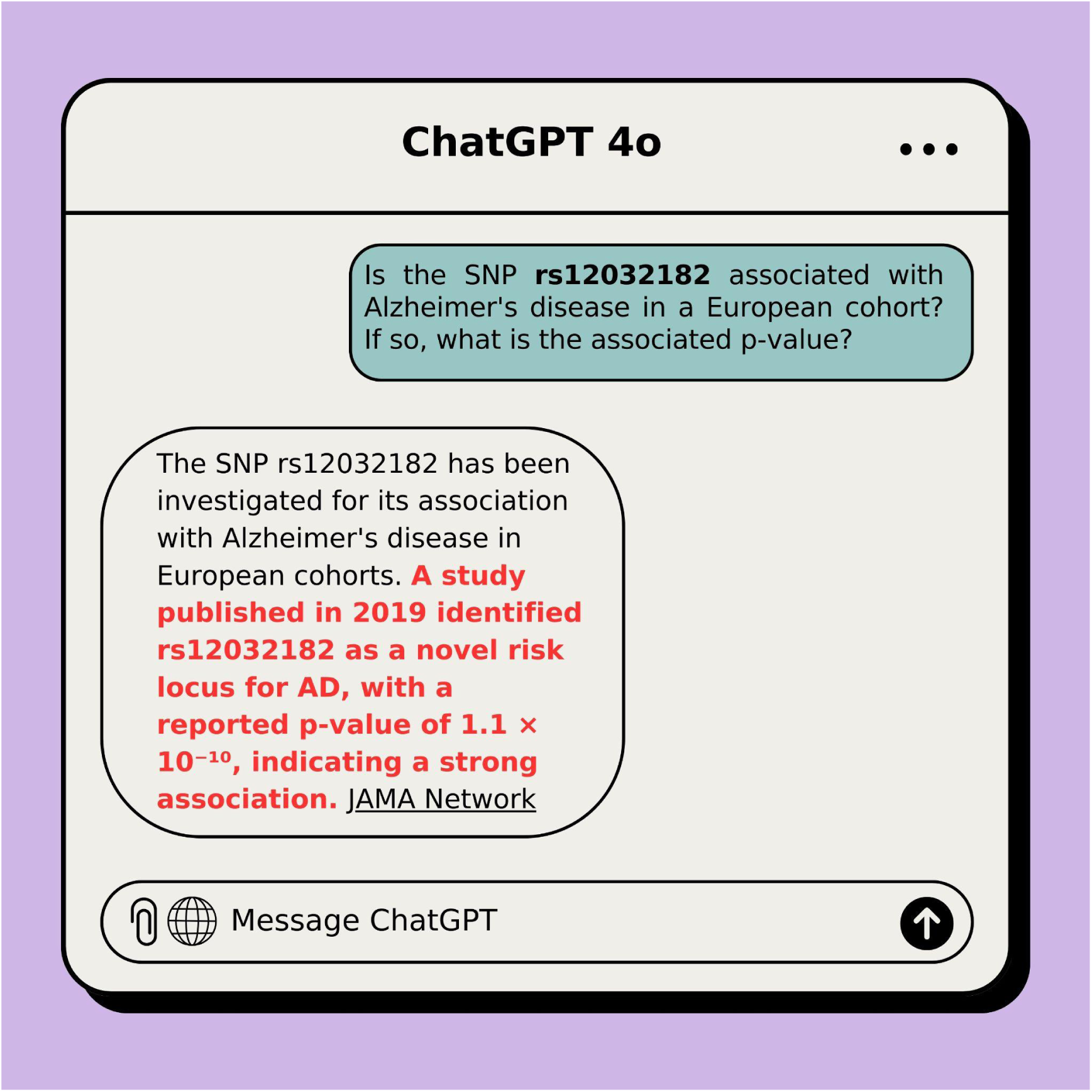
GPT-4o struggling to answer a query from CARDBiomedBench involving p-values. Highlighted in red are specific failures such as: providing a hallucinated p-value. This example highlights the limitations of current LLMs in handling specialized, data-intensive queries in the field of biological research, underscoring the need for domain-specific adaptation.

### Template Question Criteria Details

While the 80 original questions were unique when viewed in isolation, many had structural similarities when transitioned into the context of templating. For example, the original questions:

A. “*When was Quazepam assigned a United States Adopted Name (USAN) and approved for use by the FDA?”*
B. *“When was Sonidegib Phosphate assigned a United States assigned name (USAN)?”*

These questions would be considered unique on their own, however, in a template setting they would provide redundant information.

Throughout the process of selecting potential template questions, we verified that the distribution of biological categories and reasoning skills required to answer them closely reflected that of the full set, ensuring that the findings on the augmented questions were representative of the original seed questions.

Templating questions were also selected by their ability to be adapted to an automated process while maintaining accurate responses. Their structure allowed us to generate accurate responses using Python scripts by substituting variables like drug and gene names and filtering for biological logic, as demonstrated in Figure 1. This distinction in the selection process was particularly important to ensure the accuracy of our benchmark as more complex questions need a more comprehensive biological perspective that a python script can not provide.

For example:

*“Which morphinan scaffold derived medications have been modified for extended-release (ER) or sustained-release (SR) using Polistirex?”* relies on a domain expert’s knowledge of drug chemistry and categorization, which when expanded to a template question, becomes convoluted and risks comprehensiveness if answered by a script alone.

Some template questions were modified slightly for clarity. For instance, a template might request the genomic location for a single SNP instead of two, as in the original version.

**Figure S3:**
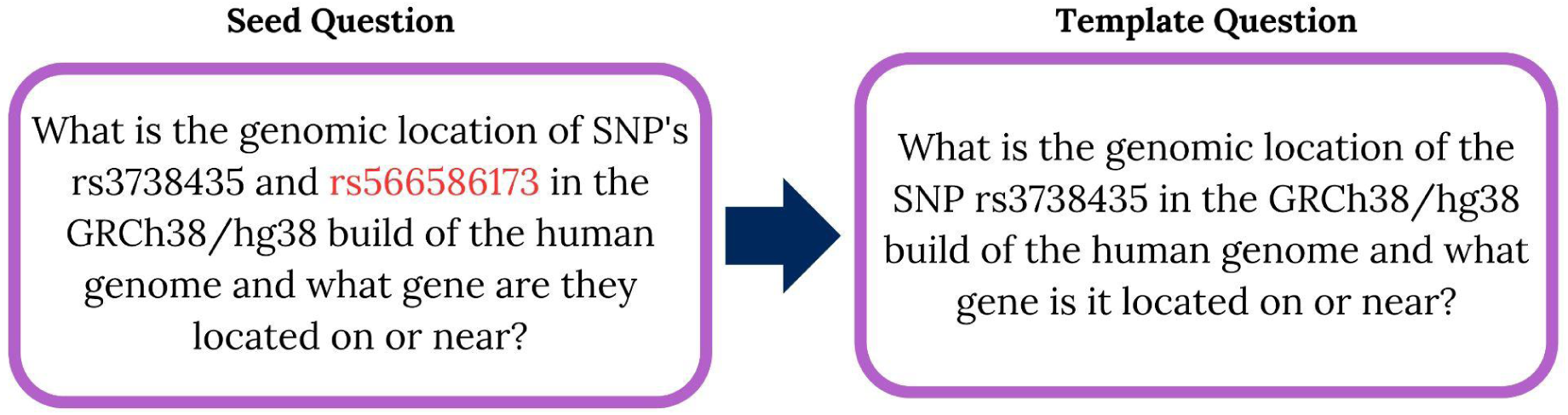
Example of refinement from a seed to a template question. The seed question requests the genomic location of two SNP’s while the template question is focused to only request one.

### Template Question Sampling Method

Since many questions produced responses that could be classified as either “Yes” or “No”, we used this to create a well distributed dataset. For questions that naturally produced less than 2,000 rows, we used all available data. For questions with over 2,000 rows, we adjusted sampling to maintain a ratio of ¾ “Yes” and ¼ “No” responses. If there were less than 1,500 “Yes” rows, we kept all “Yes” and randomly sampled the remaining “No” rows to reach 2,000. If the “Yes” rows exceeded 1,500, we took a random sample of 1,500 “Yes” and 500 “No” responses.

For running the experiments detailed in this paper, we created the “test” set by randomly sampling 270 questions from each template question where available. In the case where there were less than 270, all were included. This resulted in a “test” set of ∼10k examples.

### Specification and cost used for running models

The open-source models were run using the HuggingFace Transformers library on NIH’s BioWulf HPC at the NIH, Bethesda, MD (http://biowulf.nih.gov) which has 76 A100 nodes, each with 32 x 2.8 GHz (AMD Epyc 7543p), hyperthreading enabled, 256 MB level 3 cache, 4 x NVIDIA A100 GPUs (80 GB VRAM, 6912 cores, 432 Tensor cores), NVLINK among plenty of other computational resources. The approximate total GPU inference runtime for these experiments was 182 GPU hours in order to run the open-source models on our in-house GPU servers. The private sourced models are varying in costs/token, a breakdown of the incurred cost is shown in a table below. All models were run with their latest versions in September of 2024.

**Table S4:**
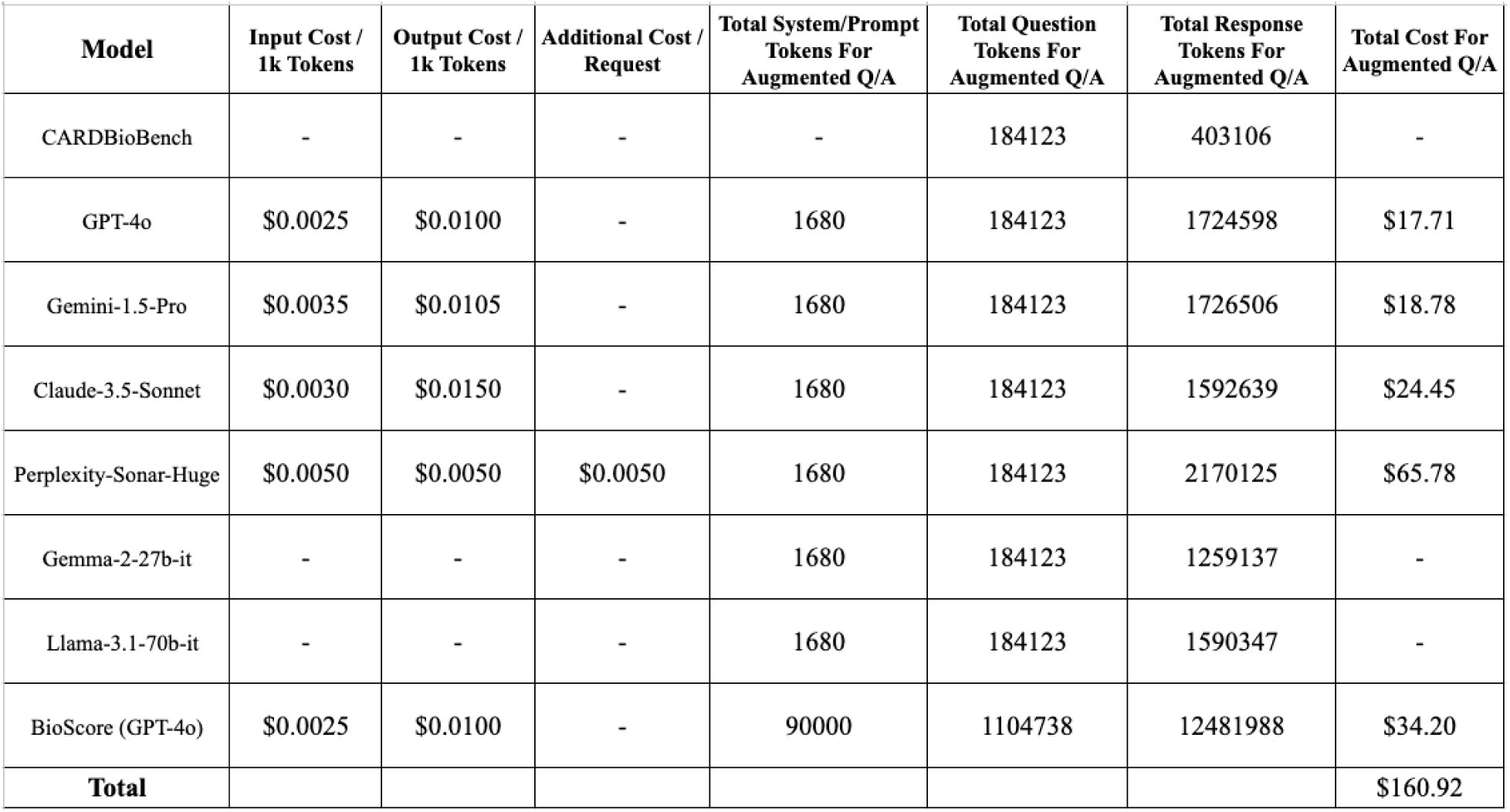
Cost breakdown of collecting responses and grading them via BioScore for our experiments. Each model has varying costs per token and number of tokens it responds with so cost is broken down by model.

We selected model hyperparameters in order to create a fair evaluation framework. Temperature was set to zero to get deterministic responses. A maximum token limit of 1024, as this was just over the benchmark answers max token count. In accordance with the known power of prompt engineering, we included a small system prompt to usher the model to respond to the questions a certain way. This was to encourage responses that aligned with the biomedical semantic space as well as give the opportunity to abstain to answer.

**Figure S5:**
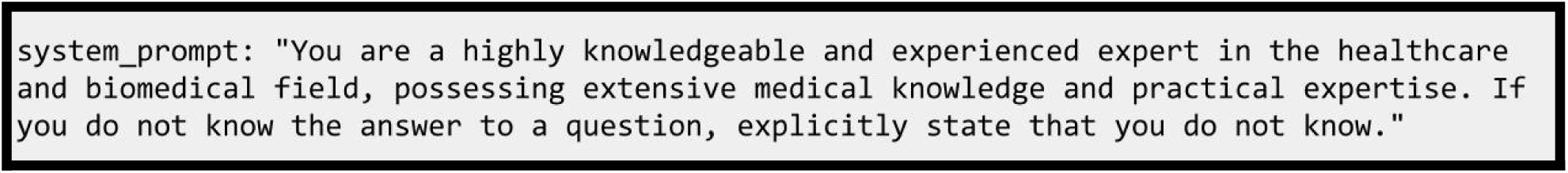
The complete system prompt given to each LLM along with the question, asking explicitly to abstain when they are unsure.

Implementation Details for BioScore

The BioScore template prompt is written below and filled in with the appropriate question, gold standard response, and predicted response. This prompt is sent to the GPT-4o API and the grades parsed from the API response. These are checked for consistency with the rubric’s scoring mechanism for a valid number. As described above, in the case of abstention the response is assigned a score of -1. These abstained questions are not included in the final BioScore, as they are counted up separately to determine the AR. Similar hyperparameters to the model responses above were used: temperature set to 0, maximum token count of 1024, and a similar system prompt without the instructions to abstain. We elected to use GPT-4o as our grading model as it is one of the most widely adopted models for evaluation and acknowledge that it may be biased towards its own responses, hence why we evaluated seven different models.

**Figure S6:**
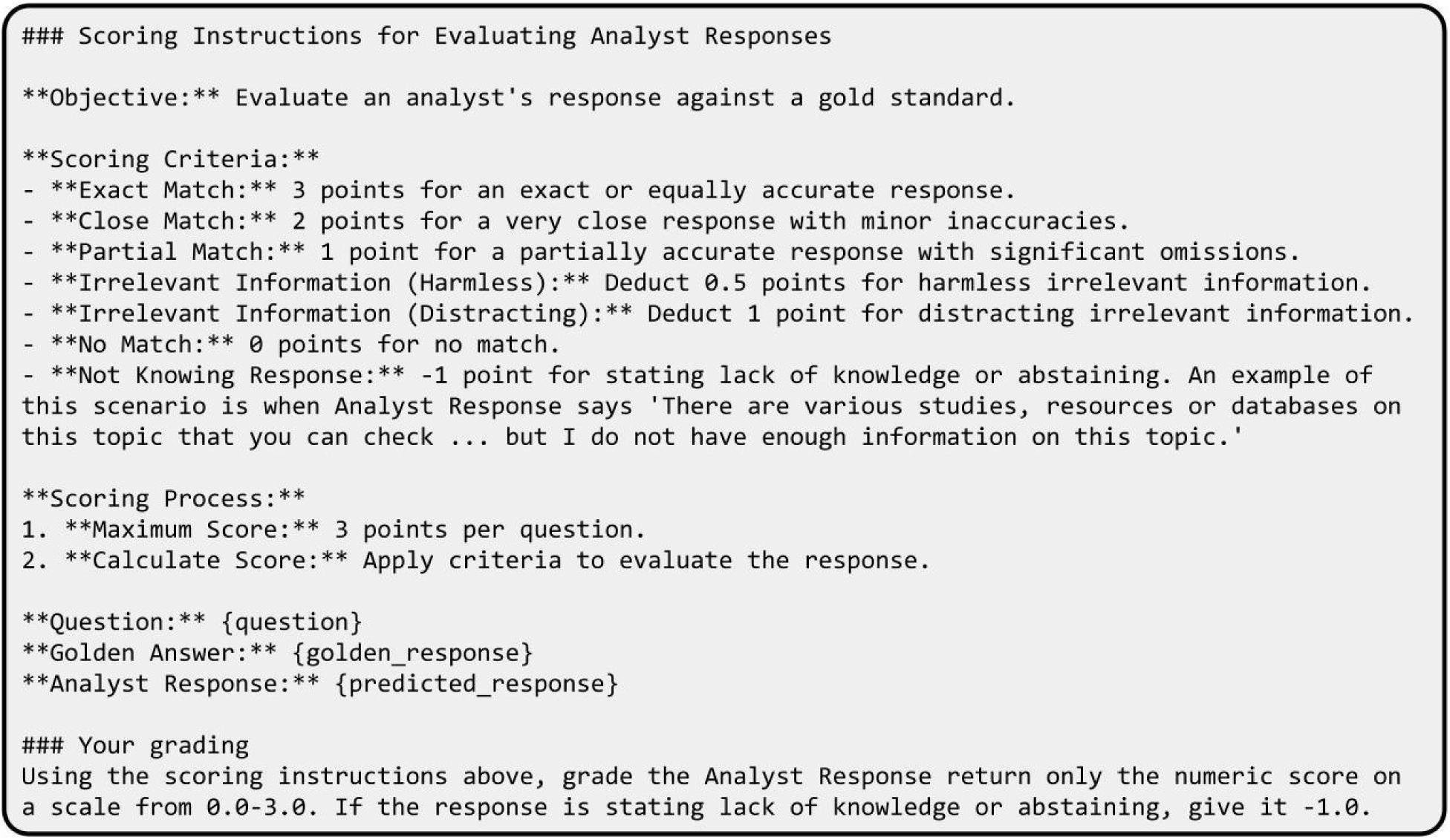
The complete BioScore grading prompt, to be filled in with appropriate question {question}, domain expert annotated “gold standard response” {golden_response}, and an LLM’s attempted answer {predicted_response}.

Our goal in creating BioScore was to design a nuanced system for assessing LLM responses that allows for various levels of correctness and relevance. We began by recognizing that not all responses would be entirely correct, so we developed a tiered scoring system to reward these varying degrees of accuracy; 3 points for exact matches, 2 points for close matches, 1 point for partial matches.

We also determined a need to account for irrelevant information by taking deductions, this too is implemented in varying degrees. For responses that contain irrelevant information that doesn’t take away from the overall message we deduct 0.5 points, to discourage unnecessary elaboration. For responses that contain irrelevant information that distracts or contradicts the overall message we deduct 1 point to reflect the negative impact on the response.

To encourage honesty we included a provision which assigned -1 points when a model reports that it doesn’t know the answer which emphasizes that it’s better to admit to a knowledge gap than to provide incorrect information.

### Error Analysis

Error analysis was conducted on the template responses to identify common failure modes of the model. This includes hallucinated responses, incomplete answers, and the generation of irrelevant information. While model abstentions are generally considered good, they were also explored in this section to better understand the models abilities. Insights from this analysis were used to refine our understanding of the model’s limitations and to suggest areas for future improvement.

**Figure S7:**
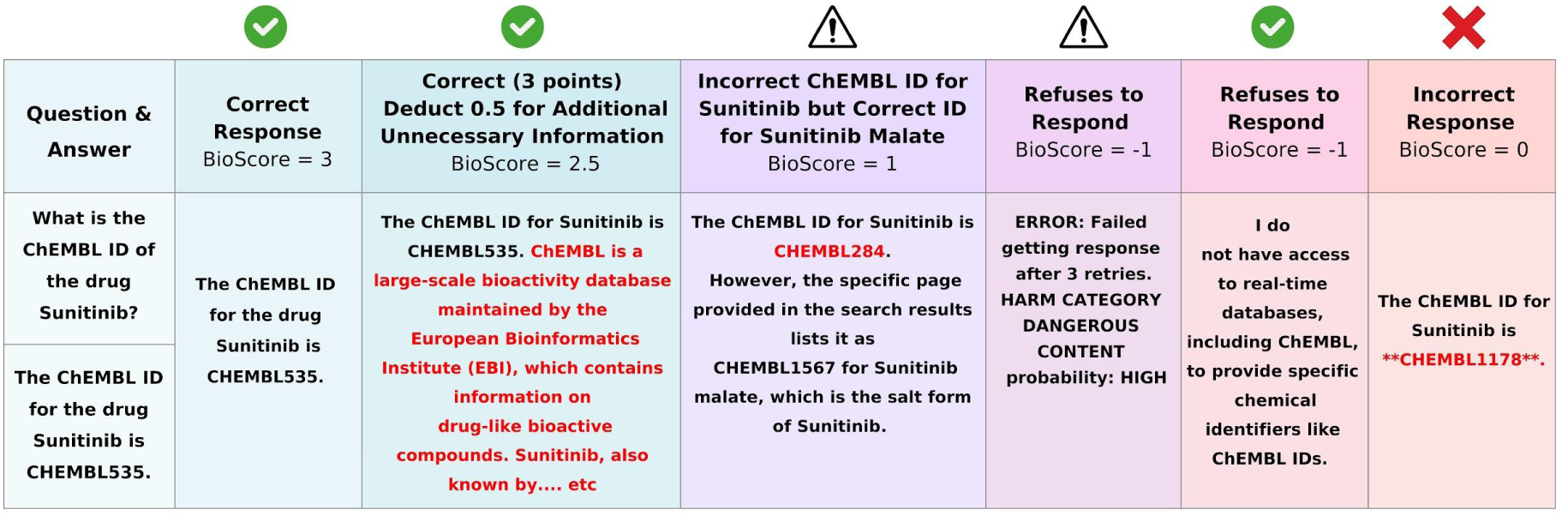
BioScore grading metric applied to the question “What is the ChEMBL ID of the drug Sunitinib?”. The first column represents the highest score, 3 points, for an exact match. In the second column, a deduction of 0.5 points is applied, yielding a BioScore of 2.5, due to unnecessary elaboration in the response. The third column illustrates an incorrect ChEMBL ID for Sunitinib but a correct ID for a related compound, resulting in a partial credit score of 1. In cases of a refusal to respond, a score of -1 is assigned, as seen in the fourth and fifth columns. Finally, an incorrect response receives a score of 0.

**Figure S8:**
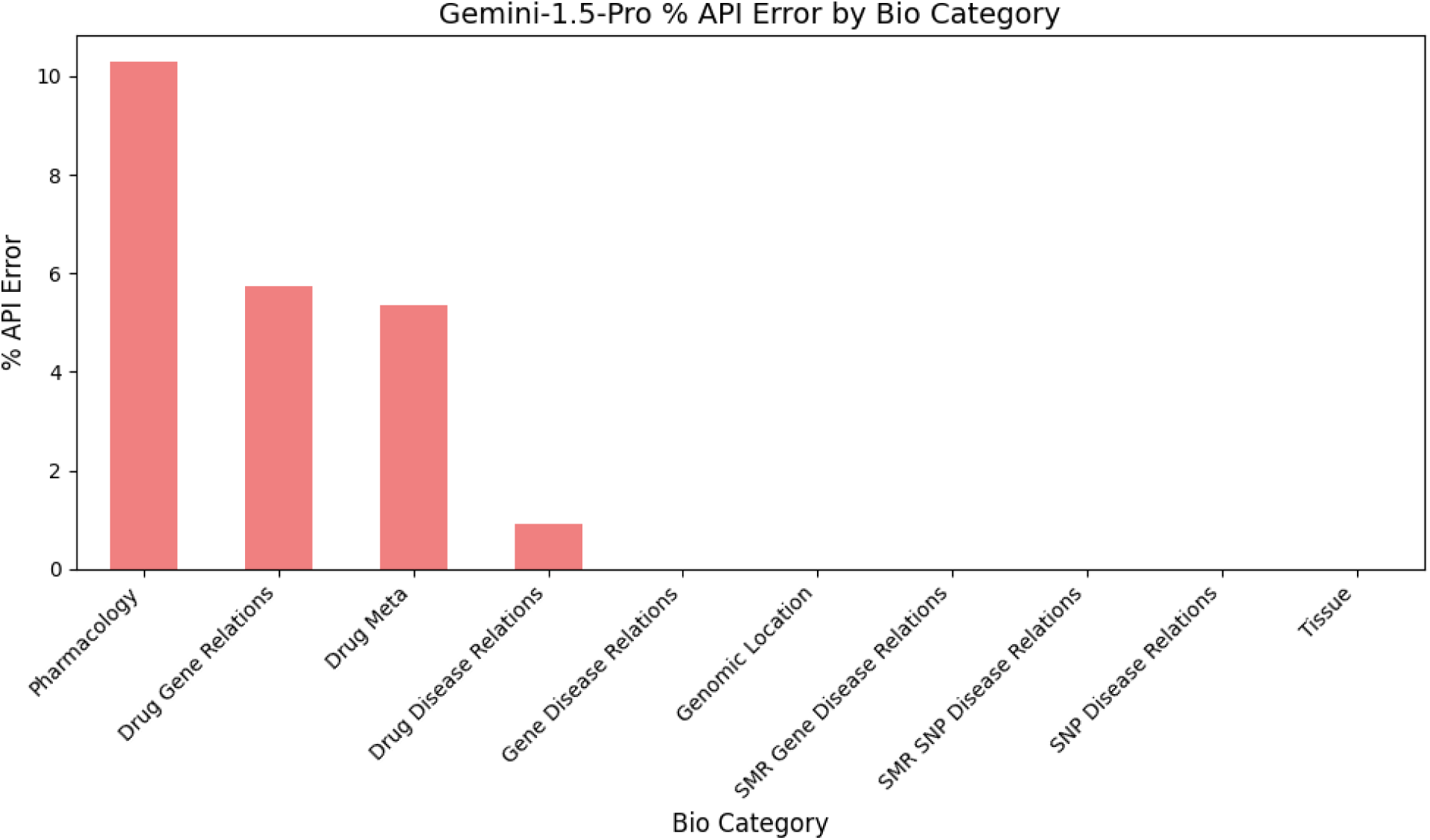
Barchart showing the percentage of Gemini API “safety errors” by Bio Category. They are a result of Gemini API’s safety filters, in particular the harm category “Dangerous Content”. Error rate can range between 0% and 100%, in the context of our Q/A lower is better as none of our questions should be deemed dangerous.

Taking a closer look at the gemini model abstentions due to API safety errors across various biological categories we can see that they occur in drug focused questions, and in particular pharmacology. These pharmacology questions aim to identify how drugs interact with biological systems, the mechanisms through which they exert effects, and specific characteristics like their molecular type or action type (e.g., as inhibitors, agonists, or binding agents). This classification is significant as pharmacology centers on understanding drug actions at both cellular and systemic levels, crucial for developing effective and safe therapeutics.

### Lexical and Semantic Scores

We also evaluated the model-generated responses using conventional lexical and semantic metrics. Lexical metrics evaluate the token overlap between model-generated and ground truth analyses. Semantic metrics evaluate the semantic similarity between the model-generated and ground truth analyses. We computed one lexical metric (BLEU),^34^ and three semantic metrics (ROUGE-1, ROUGE-L, and BERTScore).^35,36^ However, these conventional metrics of text similarity are not enough when the generated text is long and contains nuanced analysis. Figures S9 and S10 demonstrate this clearly. BioScore was able to capture the differences between a good answer and ground truth, while the traditional NLP metrics did not provide such insights. The key differences are that BioScore is able to (1) differentiate between an incorrect answer and an abstention from answering, (2) assign higher scores based on a predefined point system that awards performance according to how an expert biologist would expect an answer.

**Figure S9:**
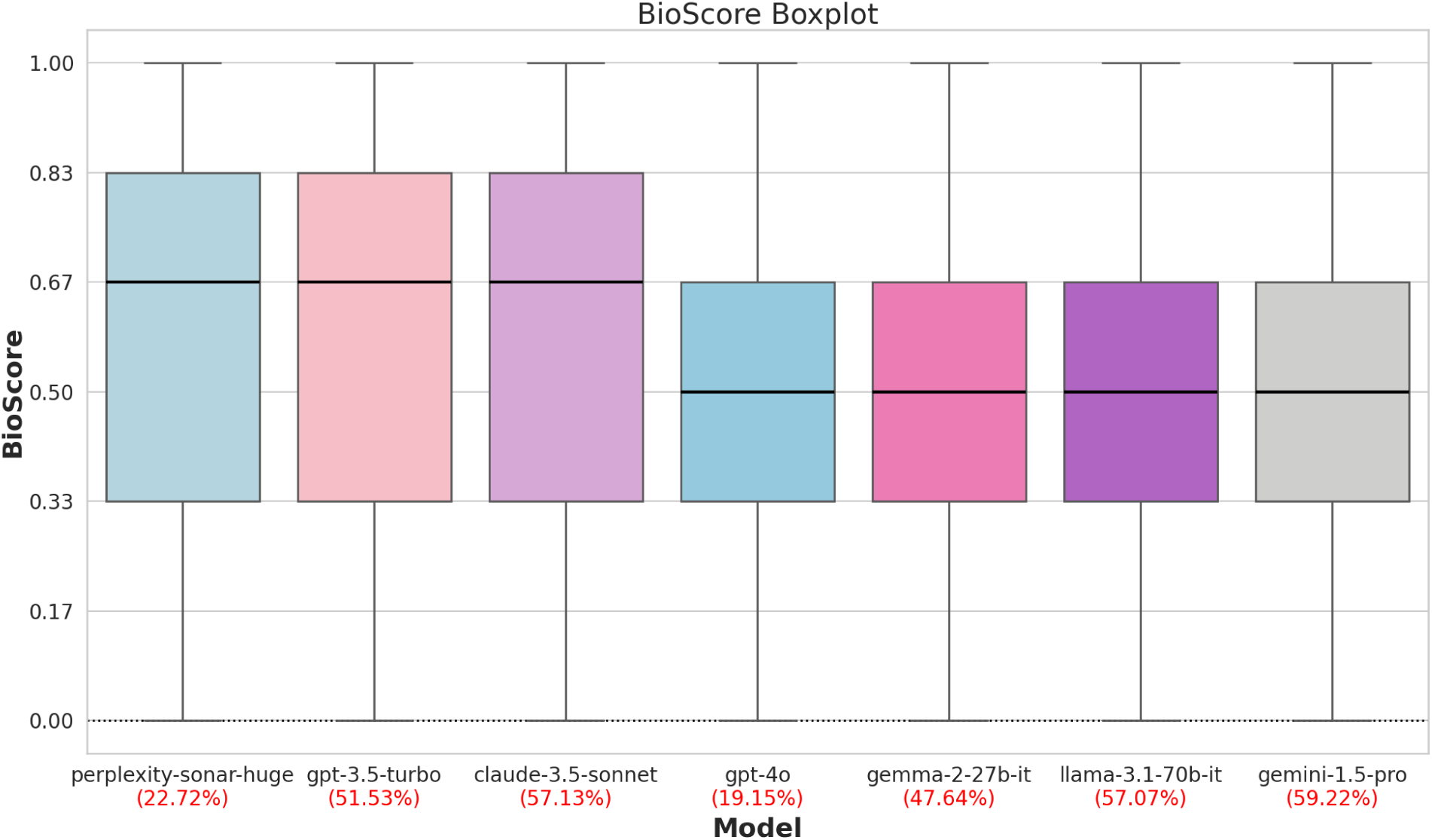
Performance of various state-of-the-art AI models on CARDBiomedBench (measured via BioScore). The Abstain Rate (AR) for each model (i.e., the ratio of the cases with the model’s self reported “I don’t know”) are also provided under each bar. A model with a higher BioScore and lower AR is more desirable. Models are sorted by decreasing median BioScore, followed by decreasing Abstain Rate (AR), and then increasing spread (interquartile range). Ranges are between 0.0 and 1.0, with higher BioScore and low AR being more desirable.

**Figure S10:**
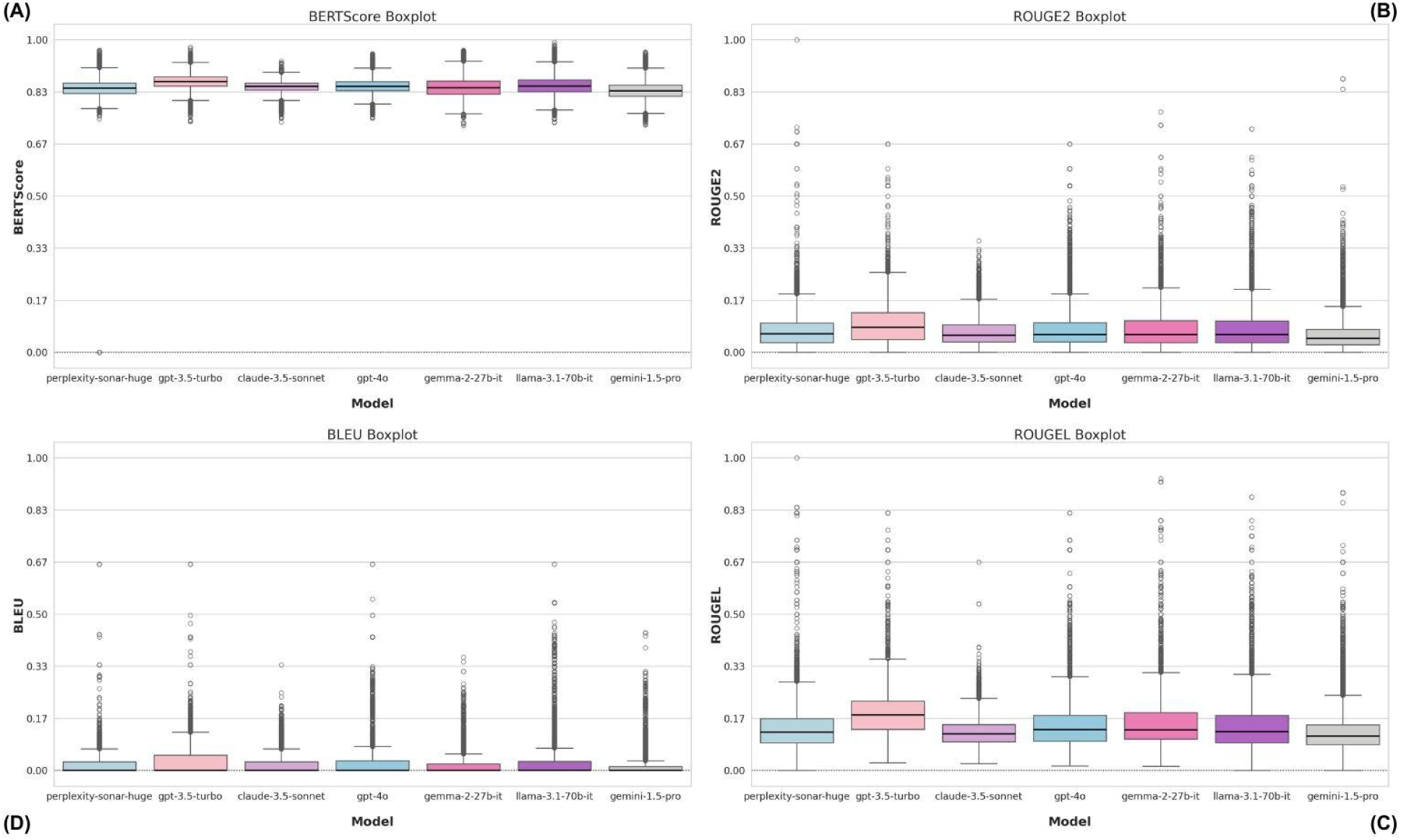
Boxplot of Performance of various state-of-the-art AI models on CARDBiomedBench (measured via traditional NLP metrics). The order of models is preserved from the Figure above. As shown, **traditional NLP metrics do not accurately capture performance on CARDBiomedBench.** This is the motivation behind our more fine-grained, rubric-based evaluation metric BioScore and accompanying AR. Ranges are between 0.0 and 1.0, with higher being more desirable.

We find that modern LLMs have significant room for improvement in the NDD domain as measured by BioScore and AR. Models all had an underwhelming performance on a subset of around 10k examples CARDBiomedBench as a whole with mean BioScore falling between 0.48 and 0.60 and AR between 0.10 and 0.60. This indicates that the models are abstaining from answering a large number of the questions and when they do respond, the quality is suffering.

**Table S11:**
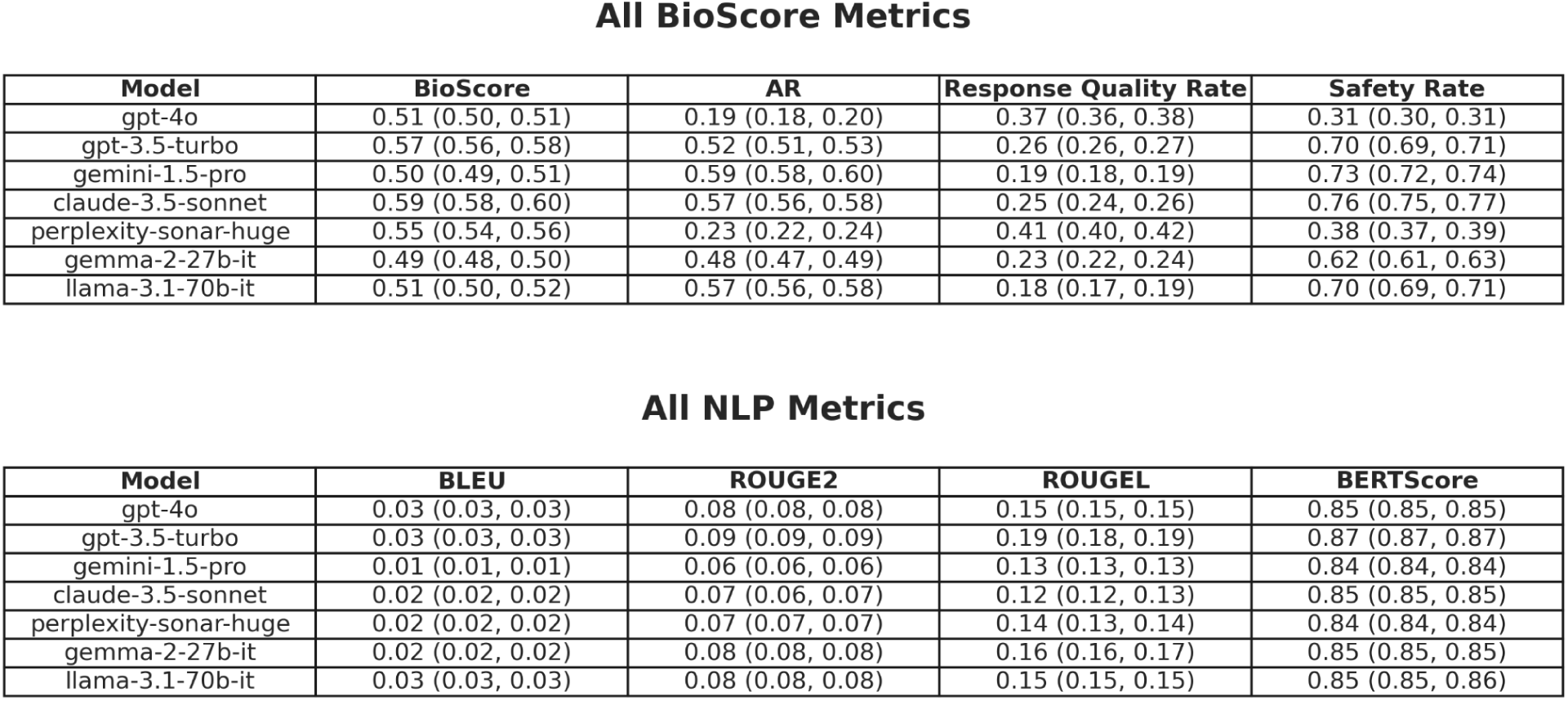
The tables report the Mean and 95% CI for each custom and NLP metric across different models. Ranges are between 0.0 and 1.0, with higher for all metrics and low AR being more desirable.

**Figure S12:**
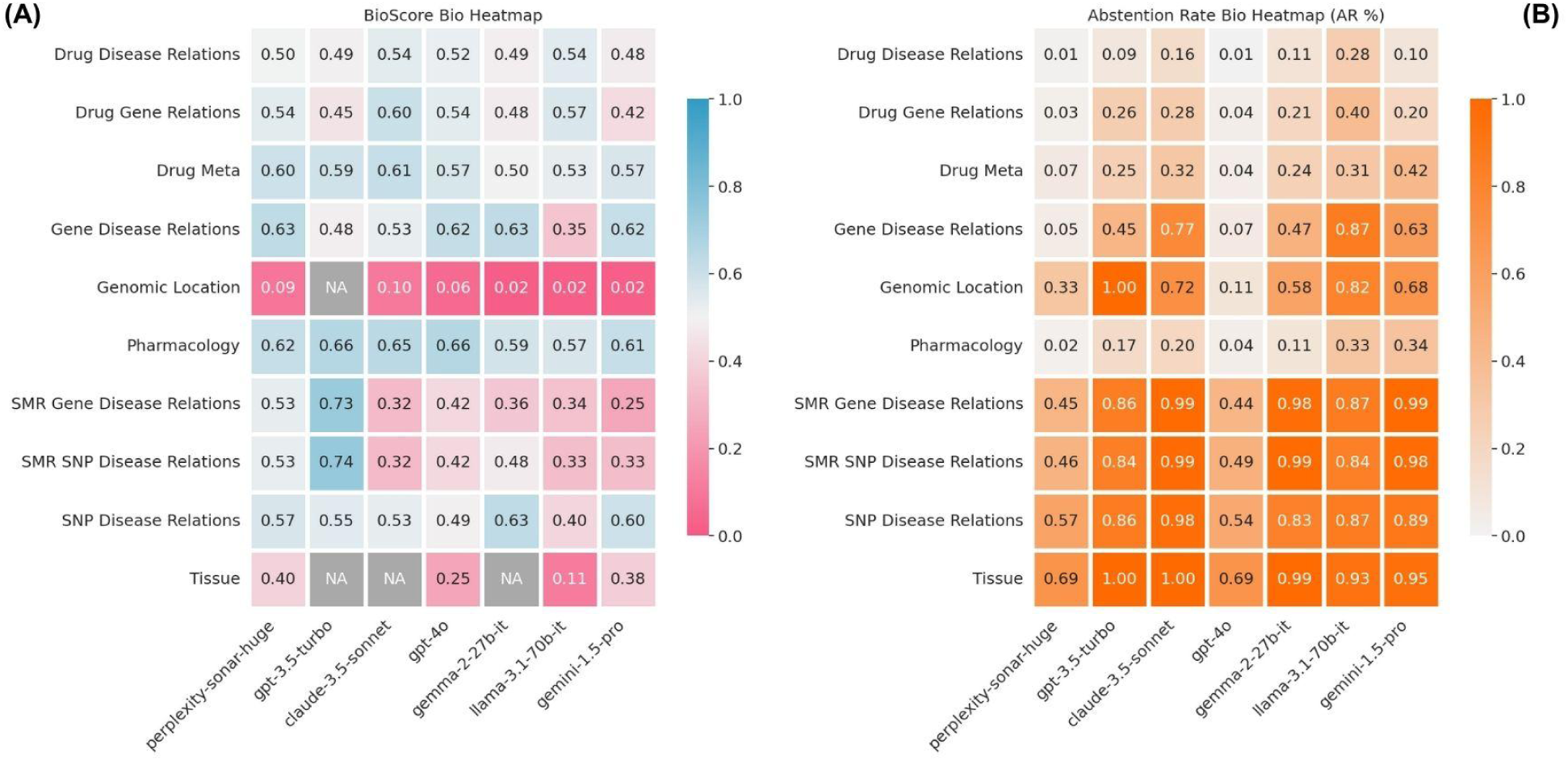
A, heatmap of mean BioScore by model (x-axis) and biological category (y-axis). B, accompanying Abstention Rates (AR). Higher BioScore (blue) and lower AR (white) are more desirable while low BioScore (red) and high AR (orange) are considered poor performance. Cells corresponding to categories with insufficient data (less than 5 responses) are displayed in dark gray and annotated with ’NA’ to denote unavailability of reliable data. Ranges are between 0.0 and 1.0, with higher BioScore and low AR being more desirable.

**Figure S13:**
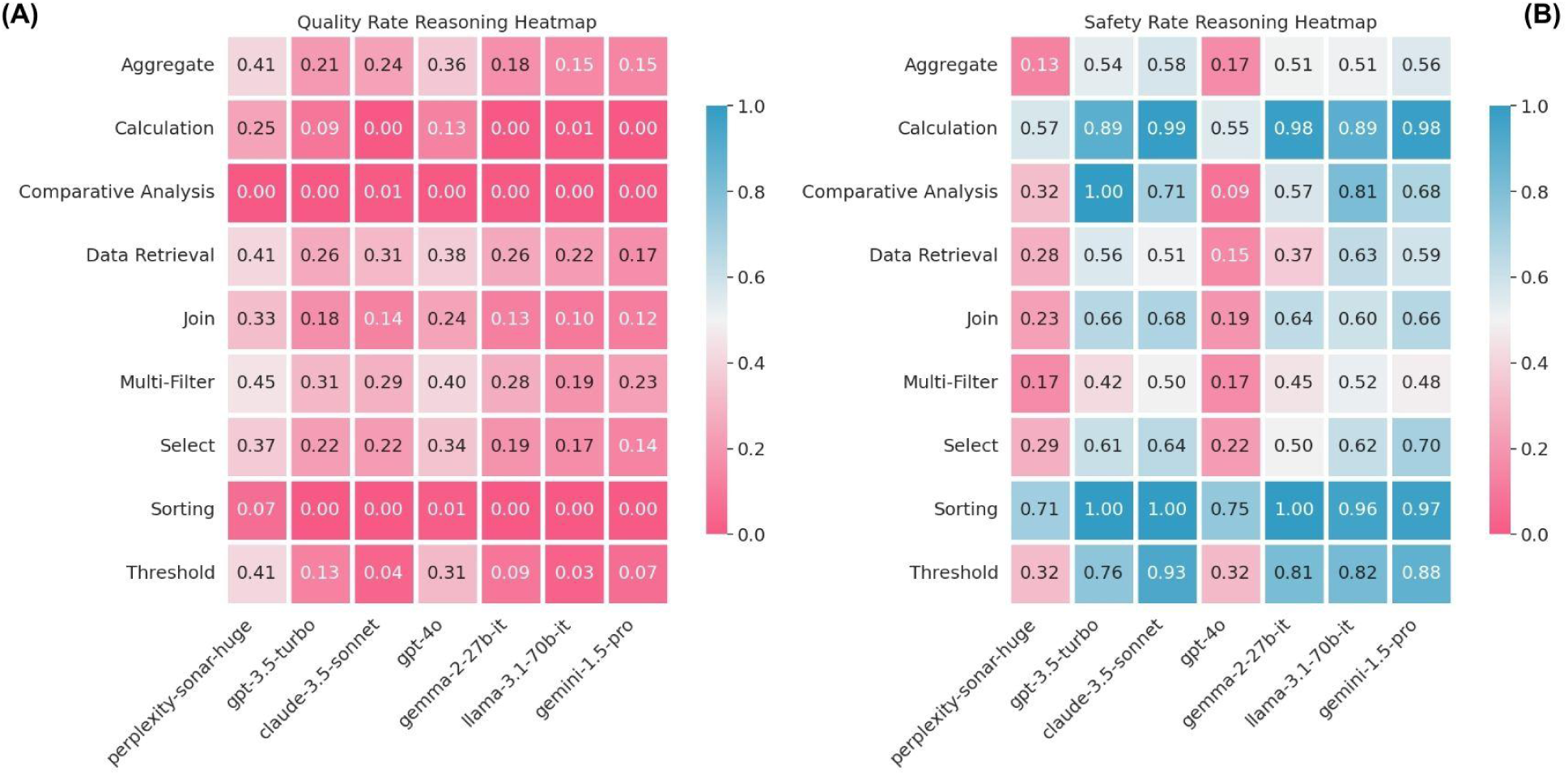
A, a heatmap of Quality Rate by model (x-axis) and reasoning category (y-axis), and B is the same heatmap Safety Rates. Higher Quality Rate and Safety Rate (blue) are more desirable while low of either (red) are considered poor performance. Ranges are between 0.0 and 1.0, with higher being more desirable.

