## Supplemental Materials for "CARDBiomedBench: A Benchmark for Evaluating Large Language Model Performance in Biomedical Research"

### Supplementary Material

#### CARDBiomedBench Statistics

| | |
|---|---|
| **# of Seed Questions** | 80 questions |
| **# of Unique Template Questions** | 40 questions |
| **# of Augmented Questions** | 68k+ questions |
| **# of Biological Categories** | 10 categories |
| **# of Reasoning Categories** | 9 categories |
| **Median Question Token Length** | 15 tokens |
| **Total Question Tokens** | 184k+ tokens |
| **Median Answer Token Length** | 34 tokens |
| **Total Answer Tokens** | 403k+ tokens |

**Table S1:** Summary of CARDBiomedBench statistics, including approximate token counts using OpenAI's tiktoken with GPT-4o as the tokenizing model.

#### Categorization of Reasoning Types

Questions in CARDBiomedBench are categorized based on complexity and operations required to retrieve data:

#### Challenging Questions

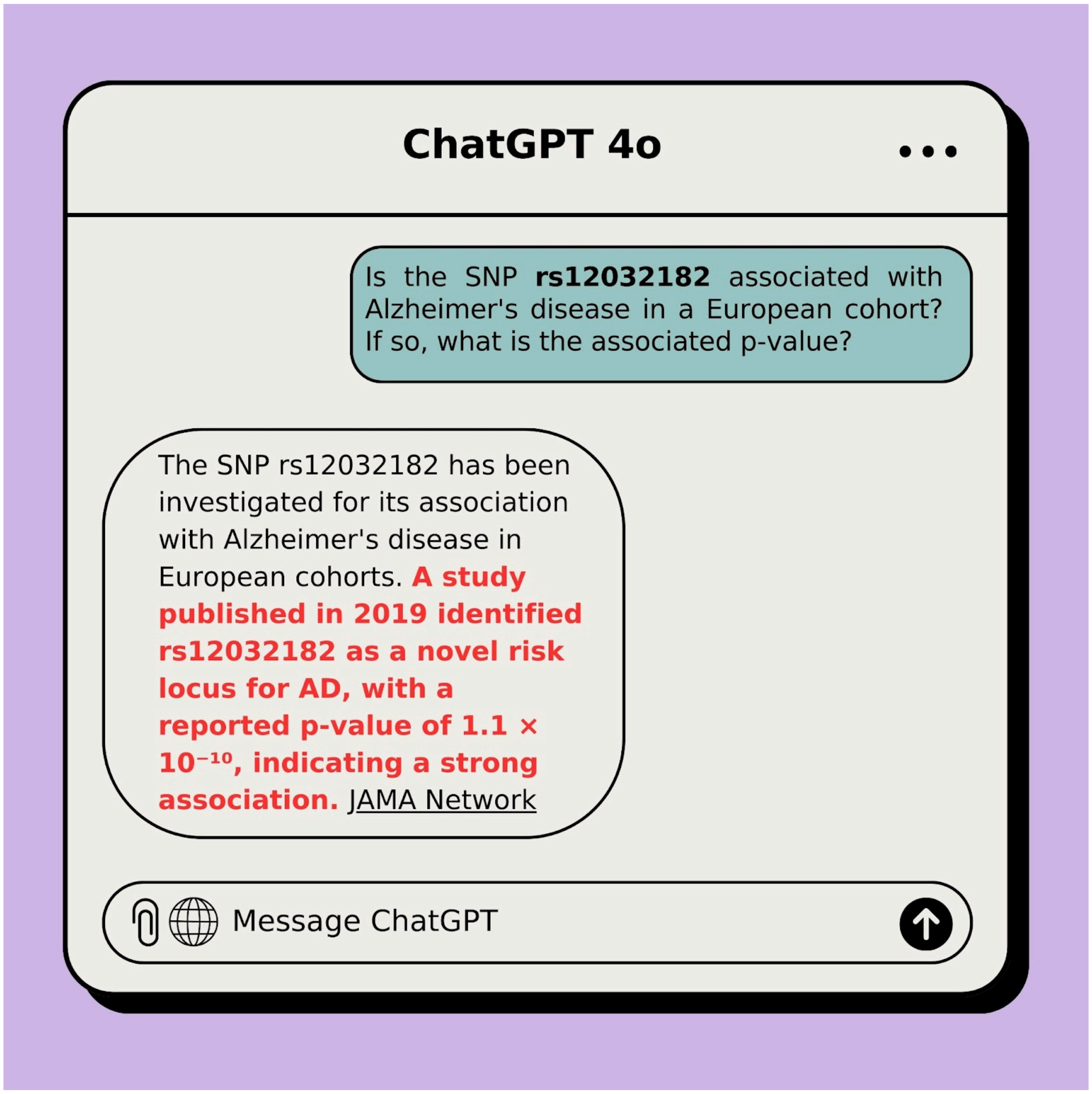


**Figure S2:** GPT-4o struggling to answer a query from CARDBiomedBench involving p-values. Highlighted in red are specific failures such as: providing a hallucinated p-value. This example highlights the limitations of current LLMs in handling specialized, data-intensive queries in the field of biological research, underscoring the need for domain-specific adaptation.

#### Template Question Criteria Details

While the 80 original questions were unique when viewed in isolation, many had structural similarities when transitioned into the context of templating. For example, the original questions:

A. "*When was Quazepam assigned a United States Adopted Name (USAN) and approved for use by the FDA?"*

*B. "When was Sonidegib Phosphate assigned a United States assigned name (USAN)?"*

These questions would be considered unique on their own, however, in a template setting they would provide redundant information.

Throughout the process of selecting potential template questions, we verified that the distribution of biological categories and reasoning skills required to answer them closely reflected that of the full set, ensuring that the findings on the augmented questions were representative of the original seed questions.

Templating questions were also selected by their ability to be adapted to an automated process while maintaining accurate responses. Their structure allowed us to generate accurate responses using Python scripts by substituting variables like drug and gene names and filtering for biological logic, as demonstrated in Figure 1. This distinction in the selection process was particularly important to ensure the accuracy of our benchmark as more complex questions need a more comprehensive biological perspective that a python script can not provide.
For example:

*"Which morphinan scaffold derived medications have been modified for extended-release (ER) or sustained-release (SR) using Polistirex?"*

relies on a domain expert's knowledge of drug chemistry and categorization, which when expanded to a template question, becomes convoluted and risks comprehensiveness if answered by a script alone.

Some template questions were modified slightly for clarity. For instance, a template might request the genomic location for a single SNP instead of two, as in the original version.

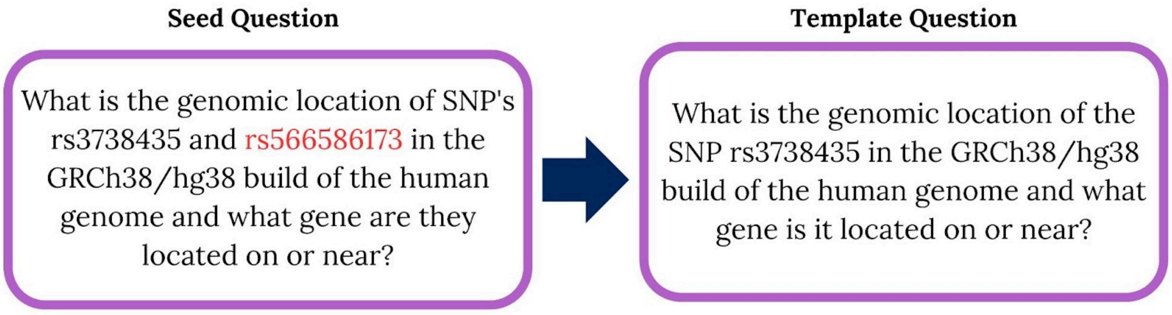


**Figure S3:** Example of refinement from a seed to a template question. The seed question requests the genomic location of two SNP's while the template question is focused to only request one.

#### Template Question Sampling Method

Since many questions produced responses that could be classified as either "Yes" or "No", we used this to create a well distributed dataset. For questions that naturally produced less than 2,000 rows, we used all available data. For questions with over 2,000 rows, we adjusted sampling to maintain a ratio of ¾ "Yes" and ¼ "No" responses. If there were less than 1,500 "Yes" rows, we kept all "Yes" and randomly sampled the remaining "No" rows to reach 2,000. If the "Yes" rows exceeded 1,500, we took a random sample of 1,500 "Yes" and 500 "No" responses.

For running the experiments detailed in this paper, we created the "test" set by randomly sampling 270 questions from each template question where available. In the case where there were less than 270, all were included. This resulted in a "test" set of ~10k examples.

#### Specification and cost used for running models

The open-source models were run using the HuggingFace Transformers library on NIH's BioWulf HPC at the NIH, Bethesda, MD (http://biowulf.nih.gov) which has 76 A100 nodes, each with 32 x 2.8 GHz (AMD Epyc 7543p), hyperthreading enabled, 256 MB level 3 cache, 4 x NVIDIA A100 GPUs (80 GB VRAM, 6912 cores, 432 Tensor cores), NVLINK among plenty of other computational resources. The approximate total GPU inference runtime for these experiments was 182 GPU hours in order to run the open-source models on our in-house GPU servers. The private sourced models are varying in costs/token, a breakdown of the incurred cost is shown in a table below. All models were run with their latest versions in September of 2024.

| Model | Input Cost / 1k Tokens | Output Cost / 1k Tokens | Additional Cost / Request | Total System/Prompt Tokens For Augmented Q/A | Total Question Tokens For Augmented Q/A | Total Response Tokens For Augmented Q/A | Total Cost For Augmented Q/A |
|---|---|---|---|---|---|---|---|
| CARDBioBench | - | - | - | - | 184123 | 403106 | - |
| GPT-4o | $0.0025 | $0.0100 | - | 1680 | 184123 | 1724598 | $17.71 |
| Gemini-1.5-Pro | $0.0035 | $0.0105 | - | 1680 | 184123 | 1726506 | $18.78 |
| Claude-3.5-Sonnet | $0.0030 | $0.0150 | - | 1680 | 184123 | 1592639 | $24.45 |
| Perplexity-Sonar-Huge | $0.0050 | $0.0050 | $0.0050 | 1680 | 184123 | 2170125 | $65.78 |
| Gemma-2-27b-it | - | - | - | 1680 | 184123 | 1259137 | - |
| Llama-3.1-70b-it | - | - | - | 1680 | 184123 | 1590347 | - |
| BioScore (GPT-4o) | $0.0025 | $0.0100 | - | 90000 | 1104738 | 12481988 | $34.20 |
| **Total** | | | | | | | $160.92 |

**Table S4:** Cost breakdown of collecting responses and grading them via BioScore for our experiments. Each model has varying costs per token and number of tokens it responds with so cost is broken down by model.

We selected model hyperparameters in order to create a fair evaluation framework. Temperature was set to zero to get deterministic responses. A maximum token limit of 1024, as this was just over the benchmark answers max token count. In accordance with the known power of prompt engineering, we included a small system prompt to usher the model to respond to the questions a certain way. This was to encourage responses that aligned with the biomedical semantic space as well as give the opportunity to abstain to answer.

```
system_prompt: "You are a highly knowledgeable and experienced expert in the healthcare
and biomedical field, possessing extensive medical knowledge and practical expertise. If
you do not know the answer to a question, explicitly state that you do not know."
```

**Figure S5:** The complete system prompt given to each LLM along with the question, asking explicitly to abstain when they are unsure.

#### Implementation Details for BioScore

The BioScore template prompt is written below and filled in with the appropriate question, gold standard response, and predicted response. This prompt is sent to the GPT-4o API and the grades parsed from the API response. These are checked for consistency with the rubric's scoring mechanism for a valid number. As described above, in the case of abstention the response is assigned a score of -1. These abstained questions are not included in the final BioScore, as they are counted up separately to determine the AR. Similar hyperparameters to the model responses above were used: temperature set to 0, maximum token count of 1024, and a similar system prompt without the instructions to abstain. We elected to use GPT-4o as our grading model as it is one of the most widely adopted models for evaluation and acknowledge that it may be biased towards its own responses, hence why we evaluated seven different models.

```
### Scoring Instructions for Evaluating Analyst Responses

**Objective:** Evaluate an analyst's response against a gold standard.

**Scoring Criteria:**
- **Exact Match:** 3 points for an exact or equally accurate response.
- **Close Match:** 2 points for a very close response with minor inaccuracies.
- **Partial Match:** 1 point for a partially accurate response with significant omissions.
- **Irrelevant Information (Harmless):** Deduct 0.5 points for harmless irrelevant information.
- **Irrelevant Information (Distracting):** Deduct 1 point for distracting irrelevant information.
- **No Match:** 0 points for no match.
- **Not Knowing Response:** -1 point for stating lack of knowledge or abstaining. An example of
this scenario is when Analyst Response says 'There are various studies, resources or databases on
this topic that you can check ... but I do not have enough information on this topic.'

**Scoring Process:**
1. **Maximum Score:** 3 points per question.
2. **Calculate Score:** Apply criteria to evaluate the response.

**Question:** {question}
**Golden Answer:** {golden_response}
**Analyst Response:** {predicted_response}

### Your grading
Using the scoring instructions above, grade the Analyst Response return only the numeric score on
a scale from 0.0-3.0. If the response is stating lack of knowledge or abstaining, give it -1.0.
```

**Figure S6:** The complete BioScore grading prompt, to be filled in with appropriate question {question}, domain expert annotated "gold standard response" {golden_response}, and an LLM's attempted answer {predicted_response}.

Our goal in creating BioScore was to design a nuanced system for assessing LLM responses that allows for various levels of correctness and relevance. We began by recognizing that not all responses would be entirely correct, so we developed a tiered scoring system to reward these varying degrees of accuracy; 3 points for exact matches, 2 points for close matches, 1 point for partial matches.

We also determined a need to account for irrelevant information by taking deductions, this too is implemented in varying degrees. For responses that contain irrelevant information that doesn't take away from the overall message we deduct 0.5 points, to discourage unnecessary elaboration. For responses that contain irrelevant information that distracts or contradicts the overall message we deduct 1 point to reflect the negative impact on the response.
To encourage honesty we included a provision which assigned -1 points when a model reports that it doesn't know the answer which emphasizes that it's better to admit to a knowledge gap than to provide incorrect information.

#### Error Analysis

Error analysis was conducted on the template responses to identify common failure modes of the model. This includes hallucinated responses, incomplete answers, and the generation of irrelevant information. While model abstentions are generally considered good, they were also explored in this section to better understand the models abilities. Insights from this analysis were used to refine our understanding of the model's limitations and to suggest areas for future improvement.

| Question & Answer | Correct Response BioScore = 3 | Correct (3 points) Deduct 0.5 for Additional Unnecessary Information BioScore = 2.5 | Incorrect ChEMBL ID for Sunitinib but Correct ID for Sunitinib Malate BioScore = 1 | Refuses to Respond BioScore = -1 | Refuses to Respond BioScore = -1 | Incorrect Response BioScore = 0 |
|---|---|---|---|---|---|---|
| What is the ChEMBL ID of the drug Sunitinib?<br>The ChEMBL ID for the drug Sunitinib is CHEMBL535. | The ChEMBL ID for the drug Sunitinib is CHEMBL535. | The ChEMBL ID for Sunitinib is CHEMBL535. ChEMBL is a large-scale bioactivity database maintained by the European Bioinformatics Institute (EBI), which contains information on drug-like bioactive compounds. Sunitinib, also known by.... etc | The ChEMBL ID for Sunitinib is CHEMBL284. However, the specific page provided in the search results lists it as CHEMBL1567 for Sunitinib malate, which is the salt form of Sunitinib. | ERROR: Failed getting response after 3 retries. HARM CATEGORY DANGEROUS CONTENT probability: HIGH | I do not have access to real-time databases, including ChEMBL, to provide specific chemical identifiers like ChEMBL IDs. | The ChEMBL ID for Sunitinib is **CHEMBL1178**. |

**Figure S7:** BioScore grading metric applied to the question "What is the ChEMBL ID of the drug Sunitinib?". The first column represents the highest score, 3 points, for an exact match. In the second column, a deduction of 0.5 points is applied, yielding a BioScore of 2.5, due to unnecessary elaboration in the response. The third column illustrates an incorrect ChEMBL ID for Sunitinib but a correct ID for a related compound, resulting in a partial credit score of 1. In cases of a refusal to respond, a score of -1 is assigned, as seen in the fourth and fifth columns. Finally, an incorrect response receives a score of 0.

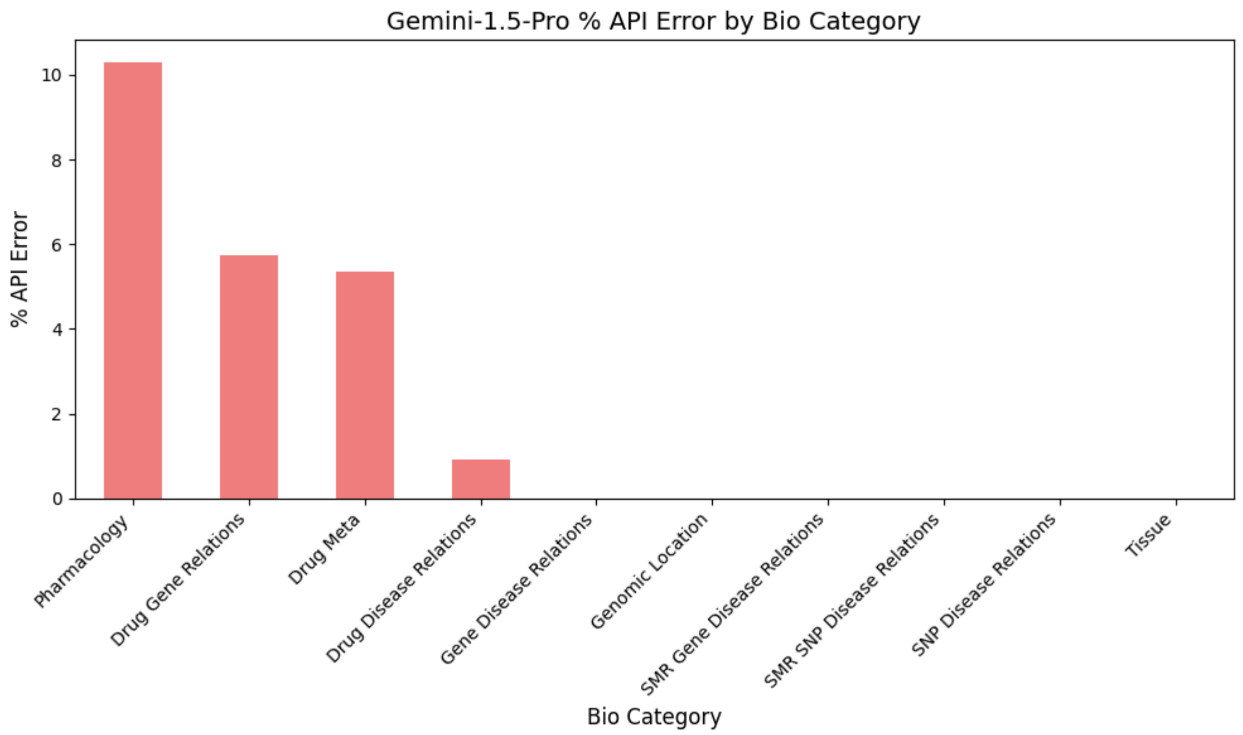


**Figure S8:** Barchart showing the percentage of Gemini API "safety errors" by Bio Category. They are a result of Gemini API's safety filters, in particular the harm category "Dangerous Content". Error rate can range between 0% and 100%, in the context of our Q/A lower is better as none of our questions should be deemed dangerous.

Taking a closer look at the gemini model abstentions due to API safety errors across various biological categories we can see that they occur in drug focused questions, and in particular pharmacology. These pharmacology questions aim to identify how drugs interact with biological systems, the mechanisms through which they exert effects, and specific characteristics like their molecular type or action type (e.g., as inhibitors, agonists, or binding agents). This classification is significant as pharmacology centers on understanding drug actions at both cellular and systemic levels, crucial for developing effective and safe therapeutics.

#### Lexical and Semantic Scores

We also evaluated the model-generated responses using conventional lexical and semantic metrics. Lexical metrics evaluate the token overlap between model-generated and ground truth analyses. Semantic metrics evaluate the semantic similarity between the model-generated and ground truth analyses. We computed one lexical metric (BLEU),[34] and three semantic metrics (ROUGE-1, ROUGE-L, and BERTScore).[35,36] However, these conventional metrics of text similarity are not enough when the generated text is long and contains nuanced analysis. Figures S9 and S10 demonstrate this clearly. BioScore was able to capture the differences between a good answer and ground truth, while the traditional NLP metrics did not provide such insights. The key differences are that BioScore is able to (1) differentiate between an incorrect answer and an abstention from answering, (2) assign higher scores based on a predefined point system that awards performance according to how an expert biologist would expect an answer.

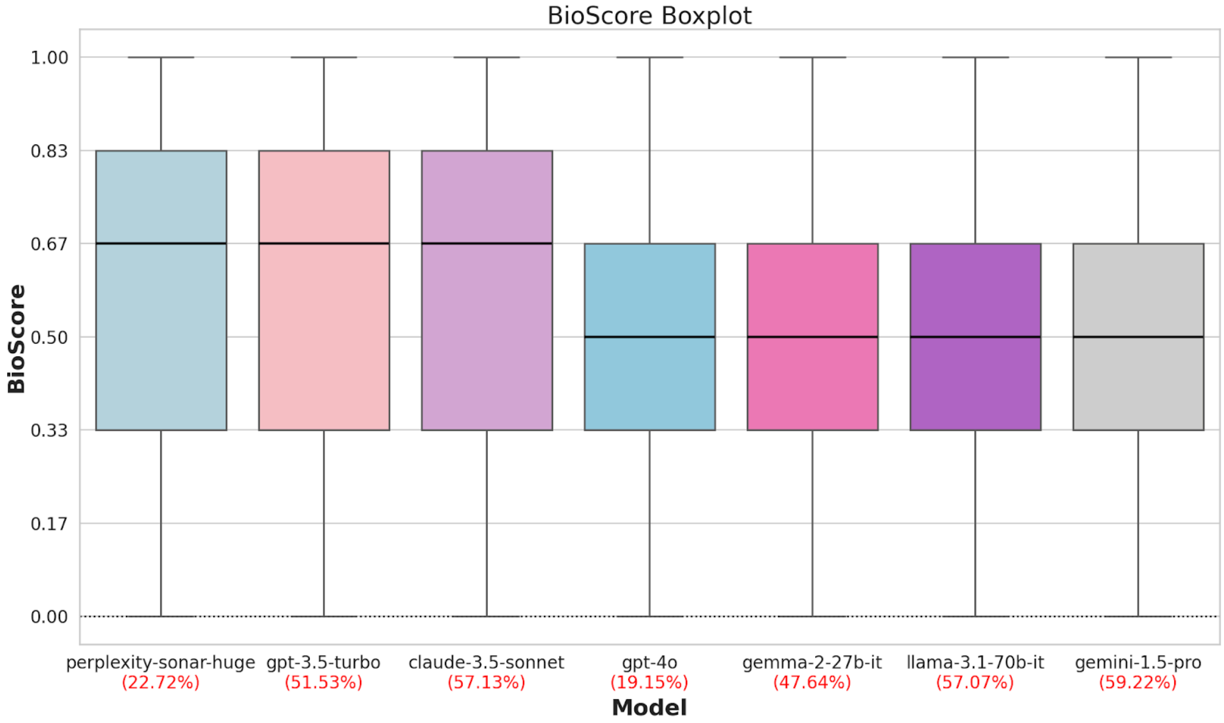


**Figure S9:** Performance of various state-of-the-art AI models on CARDBiomedBench (measured via BioScore). The Abstain Rate (AR) for each model (i.e., the ratio of the cases with the model's self reported "I don't know") are also provided under each bar. A model with a higher BioScore and lower AR is more desirable. Models are sorted by decreasing median BioScore, followed by decreasing Abstain Rate (AR), and then increasing spread (interquartile range). Ranges are between 0.0 and 1.0, with higher BioScore and low AR being more desirable.

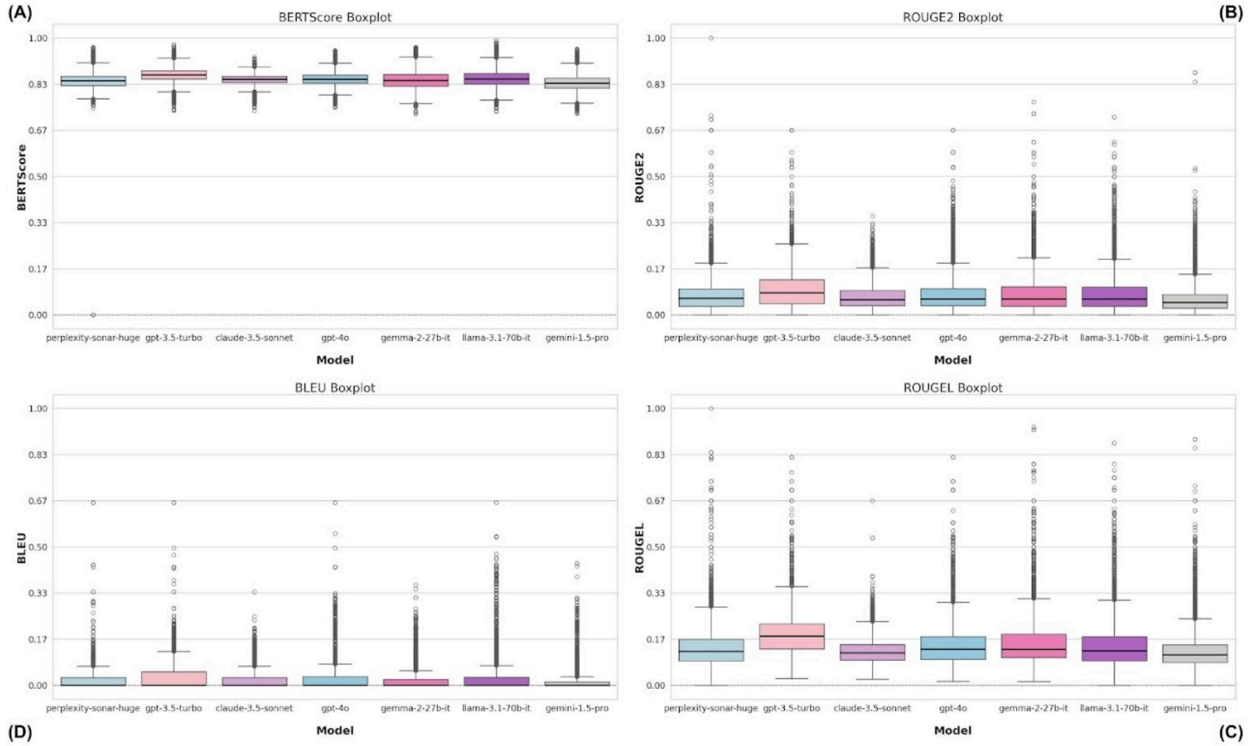


**Figure S10:** Boxplot of Performance of various state-of-the-art AI models on CARDBiomedBench (measured via traditional NLP metrics). The order of models is preserved from the Figure above. As shown, **traditional NLP metrics do not accurately capture performance on CARDBiomedBench.** This is the motivation behind our more fine-grained, rubric-based evaluation metric BioScore and accompanying AR. Ranges are between 0.0 and 1.0, with higher being more desirable.

We find that modern LLMs have significant room for improvement in the NDD domain as measured by BioScore and AR. Models all had an underwhelming performance on a subset of around 10k examples CARDBiomedBench as a whole with mean BioScore falling between 0.48 and 0.60 and AR between 0.10 and 0.60. This indicates that the models are abstaining from answering a large number of the questions and when they do respond, the quality is suffering.

#### All BioScore Metrics

| Model | BioScore | AR | Response Quality Rate | Safety Rate |
|---|---|---|---|---|
| gpt-4o | 0.51 (0.50, 0.51) | 0.19 (0.18, 0.20) | 0.37 (0.36, 0.38) | 0.31 (0.30, 0.31) |
| gpt-3.5-turbo | 0.57 (0.56, 0.58) | 0.52 (0.51, 0.53) | 0.26 (0.26, 0.27) | 0.70 (0.69, 0.71) |
| gemini-1.5-pro | 0.50 (0.49, 0.51) | 0.59 (0.58, 0.60) | 0.19 (0.18, 0.19) | 0.73 (0.72, 0.74) |
| claude-3.5-sonnet | 0.59 (0.58, 0.60) | 0.57 (0.56, 0.58) | 0.25 (0.24, 0.26) | 0.76 (0.75, 0.77) |
| perplexity-sonar-huge | 0.55 (0.54, 0.56) | 0.23 (0.22, 0.24) | 0.41 (0.40, 0.42) | 0.38 (0.37, 0.39) |
| gemma-2-27b-it | 0.49 (0.48, 0.50) | 0.48 (0.47, 0.49) | 0.23 (0.22, 0.24) | 0.62 (0.61, 0.63) |
| llama-3.1-70b-it | 0.51 (0.50, 0.52) | 0.57 (0.56, 0.58) | 0.18 (0.17, 0.19) | 0.70 (0.69, 0.71) |

#### All NLP Metrics

| Model | BLEU | ROUGE2 | ROUGEL | BERTScore |
|---|---|---|---|---|
| gpt-4o | 0.03 (0.03, 0.03) | 0.08 (0.08, 0.08) | 0.15 (0.15, 0.15) | 0.85 (0.85, 0.85) |
| gpt-3.5-turbo | 0.03 (0.03, 0.03) | 0.09 (0.09, 0.09) | 0.19 (0.18, 0.19) | 0.87 (0.87, 0.87) |
| gemini-1.5-pro | 0.01 (0.01, 0.01) | 0.06 (0.06, 0.06) | 0.13 (0.13, 0.13) | 0.84 (0.84, 0.84) |
| claude-3.5-sonnet | 0.02 (0.02, 0.02) | 0.07 (0.06, 0.07) | 0.12 (0.12, 0.13) | 0.85 (0.85, 0.85) |
| perplexity-sonar-huge | 0.02 (0.02, 0.02) | 0.07 (0.07, 0.07) | 0.14 (0.13, 0.14) | 0.84 (0.84, 0.84) |
| gemma-2-27b-it | 0.02 (0.02, 0.02) | 0.08 (0.08, 0.08) | 0.16 (0.16, 0.17) | 0.85 (0.85, 0.85) |
| llama-3.1-70b-it | 0.03 (0.03, 0.03) | 0.08 (0.08, 0.08) | 0.15 (0.15, 0.15) | 0.85 (0.85, 0.86) |

**Table S11:** The tables report the Mean and 95% CI for each custom and NLP metric across different models. Ranges are between 0.0 and 1.0, with higher for all metrics and low AR being more desirable.

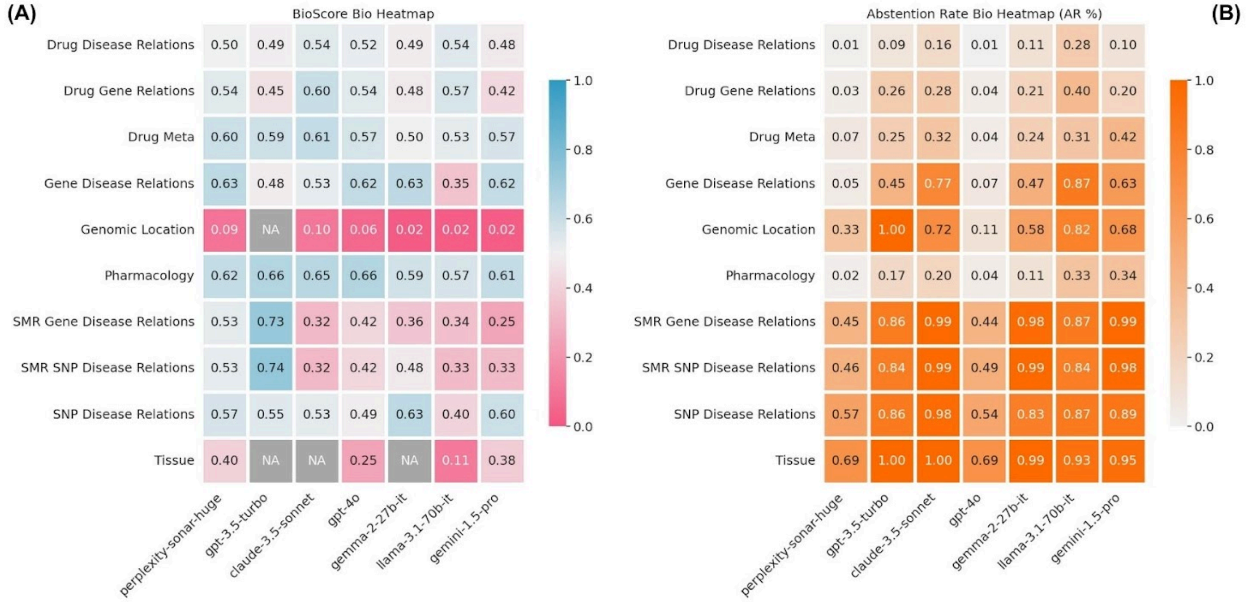


**Figure S12:** A, heatmap of mean BioScore by model (x-axis) and biological category (y-axis). B, accompanying Abstention Rates (AR). Higher BioScore (blue) and lower AR (white) are more desirable while low BioScore (red) and high AR (orange) are considered poor performance. Cells corresponding to categories with insufficient data (less than 5 responses) are displayed in dark gray and annotated with 'NA' to denote unavailability of reliable data. Ranges are between 0.0 and 1.0, with higher BioScore and low AR being more desirable.

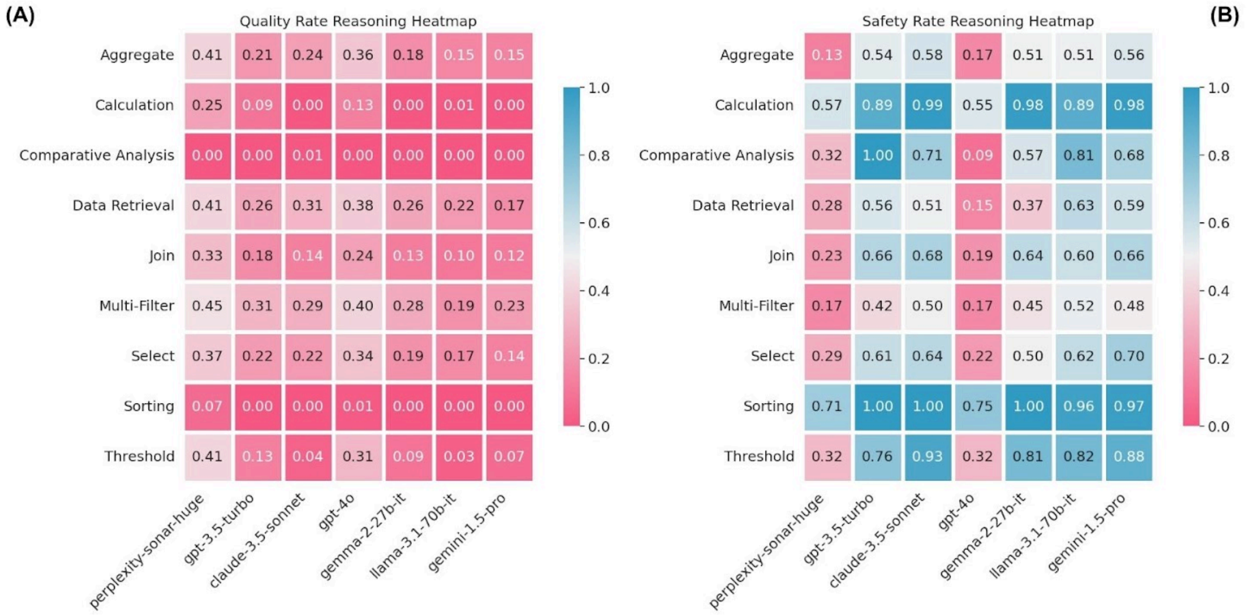


**Figure S13:** A, a heatmap of Quality Rate by model (x-axis) and reasoning category (y-axis), and B is the same heatmap Safety Rates. Higher Quality Rate and Safety Rate (blue) are more desirable while low of either (red) are considered poor performance. Ranges are between 0.0 and 1.0, with higher being more desirable.
